# Neural Circuit Mechanisms of Representation Transformation in Sequential Economic Choice

**DOI:** 10.1101/2025.08.29.673193

**Authors:** Jintao Gu, Kuntan Ni, Ben Vengerik, Xinying Cai, Sukbin Lim

## Abstract

Choices under sequential offers require the brain to encode a current offer, maintain its value, and compare it with a later offer before selecting an action. Prominent models explain choice through attractor dynamics that maintain stable, identity-preserving representations of offers and choices. However, recent recordings from monkey orbitofrontal cortex (OFC) during sequential-offer economic choices revealed strong order-based choice signals and weak identity-based signals. To identify the circuit principles underlying these representations, we trained recurrent neural networks (RNNs) to perform an order-based choice task and reproduce the observed psychometric behavior. The networks developed overlapping value representations across offer identities and dynamically reformatted the first offer across delay periods into a memory signal that could be compared with the incoming second offer. Low-dimensional latent-circuit inference revealed complementary roles for feedforward and recurrent connectivity: feedforward weights mapped different offer types onto a common value scale, whereas asymmetric recurrent interactions between sensory and decision variables generated order-based choice signals. Relationships among value coding across task phases and order-choice preference were qualitatively consistent with monkey OFC data. In a controlled comparison, RNNs trained for juice-based choice exhibited relatively stable, attractor-like representations, yet these could coexist with order-based transformations within individual networks. Together, these findings illustrate how recurrent circuits can flexibly reformat value representations across computational stages to support memory-guided decisions, linking this population-level reorganization to a specific dynamical circuit motif.

**Significance Statement:** Attractor dynamics have long provided a powerful framework for working memory and decision-making, yet recent studies across cognitive tasks suggest that neural representations can reorganize over time. Similar questions arise in sequential-offer economic choice, where stable and dynamic value-coding schemes have been linked to commodity-based and order-based decisions, respectively. Here, we used task-optimized recurrent neural networks to move from experimentally observed order-based choice signals to a mechanistic account of the circuit and neural signatures, revealing commodity-agnostic value coding and dynamic reformatting of value into a memory representation for comparison with a subsequent offer. Our work broadens theories of decision-making beyond stable attractors and provides a foundation for understanding more ethological choices that engage multiple representational formats and diverse circuit motifs.

## Introduction

Cognitive computations unfold over time, and neural representations often change with them. Rather than remaining fixed, population activity can reorganize within (Churchland et al., 2012) or across successive processing stages in sensorimotor tasks (Elsayed et al., 2016; Remington et al., 2018). In working memory tasks, reformatting of representations has been proposed to reduce interference between stored and incoming information (Spaak et al., 2017; Cavanagh et al., 2018; Parthasarathy et al., 2019; Libby and Buschman, 2021; Yang and Yu, 2025), or to prioritize behaviorally relevant items under attentional control (Panichello and Buschman, 2021; Jahn et al., 2024). Similarly, during decision-making, neural activity often evolves from encoding sensory or value information to supporting memory, comparison, and action selection, suggesting that population geometry is dynamically reshaped as computation proceeds (Aoi et al., 2020; Yoo and Hayden, 2020; Johnston et al., 2024).

The circuit mechanisms underlying this reorganization remain unclear. This issue is particularly salient in sequential-offer economic decision-making, where the brain must encode one offer, retain its value in memory, and compare it with a later offer to produce a choice (Strait et al., 2014; Azab and Hayden, 2017; Ballesta and Padoa-Schioppa, 2019; Shi et al., 2022). Classical circuit models based on attractor dynamics have provided an influential framework for understanding stable memory states and robust choice formation (Wang, 2002, 2008). Especially under simultaneous presentations of alternatives during economic choices, attractor dynamics are thought to arise from competition between neural populations or subspaces associated with each choice, defined by the spatial attributes or identities of the offers (Hunt et al., 2012; Chau et al., 2014; Rustichini and Padoa-Schioppa, 2015). However, it remains less clear how recurrent circuits generate the representational transformations observed in sequential-offer tasks, in which information must be flexibly reformatted across task phases.

Recent experimental results in sequential-offer choice tasks indicate that behavior is strongly influenced by presentation order, especially when choice reporting does not require maintenance of offer identity, pointing to computations that extend beyond attractor-based models with stable stimulus and choice representations. To investigate the circuit principles that support these computations, we trained RNNs to perform an order-based choice task. We then analyzed their activity at the levels of single neurons, population dynamics, and latent circuit structure. As a control, we also examined RNNs trained on a juice-based choice task to determine how the representational and dynamical signatures identified in the order-task networks differed from those associated with stable, identity-based choice. The order-task networks reveal how value signals are maintained and converted into choices through the structured reorganization of population activity. The model-derived relationships among coding axes and their transformation across task phases were qualitatively consistent with those observed in monkey OFC recordings, providing a mechanistic framework for understanding how recurrent circuits support flexible sequential decisions.

## Materials and Methods

### Animal Task

Two adult male rhesus monkeys (*Macaca mulatta*; monkey N, 11.5 kg; monkey F, 17.0 kg) were Involved in this study. All experimental procedures conformed to the NIH *Guide for the Care and Use of Laboratory Animals* and were approved by the Institutional Animal Care and Use Committee at NYU Shanghai. The animals were trained to choose between two juices sequentially offered in variable amounts (Fig. 1A). They were head-fixed, presented with visual stimuli on a computer monitor, and their eye positions were monitored to collect their responses. The offers were presented on either side of the screen while the monkeys fixate on the center of the screen. After Offer 2 disappeared, the monkeys saccade to one of two targets, and were rewarded with the offer shown on the side of the saccade. The juice presentation order (AB/BA) and the spatial configuration were randomized across trials.

**Figure 1.**
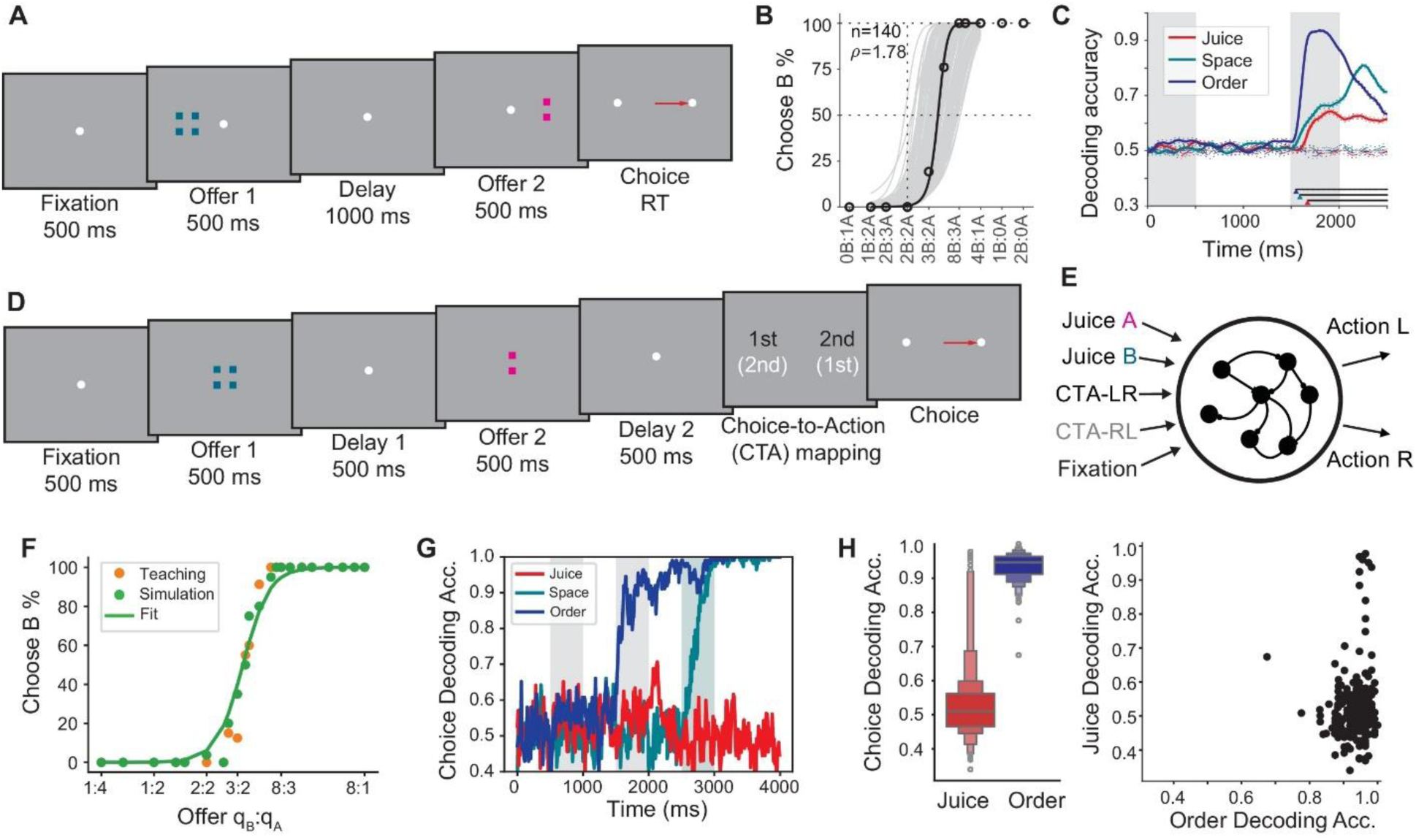
Order-based choice in experiment (A-C) and its replication in task-optimized RNNs (D-H). (A) Task schematic of monkey experiment. Colors and square numbers represent juice type and quantity. (B) (C) Decoding of choices from OFC activity across reference frames. Solid lines: mean accuracy; dotted: ± SEM; dashed: shuffle controls. (D) Modified task design for RNN training, including a two-type choice-to-action (CTA) mapping cue after the offers. (E) Network architecture of the RNN. (F,G) Choice behavior (F) and decoding accuracy (H) from the example RNN. Orange circles: random instantiation of the teaching psychometric curve; blue circles: behavior output of the example RNN; green line: fitted curve to RNN’s responses. Gray and blue shading indicate offer presentation and CTA cue, respectively. (H) Decoding accuracy of juice- and order-based choices across the RNN ensemble.

### Electrophysiology

After the animals achieved stable task performance, craniotomy surgeries were performed to open rectangular recording windows – on the left hemisphere of monkey N (AP 32:38, ML 7:17 mm, anterior limit of the corpus callosum is at A34) and at the right hemisphere of monkey F (AP 32:38, L 6:18 mm, anterior limit of the corpus callosum is at A34). Acute extracellular recordings were performed with single or double 16-channel linear arrays within or across OFC (824 neurons for monkey N and 402 neurons for monkey F). Recording sites varied across sessions. In each session, the relative value of the juices was estimated by logistic regression with a sliding window. We used SpyKING CIRCUS (Yger et al., 2018), an open-source semi-automated spike-sorting software based on a template-matching algorithm, to detect as many units as possible. This was followed by manual curation based on analyses of spike waveform features and inter-spike-interval (ISI) distributions. Only pure single units and multi-units containing one dominant unit were included in subsequent analyses. To reduce systematic firing-rate drift potentially associated with semi-automated spike sorting, the baseline activity of each trial, defined by the mean firing rate during the center fixation period, was subtracted from the firing rates at each time. Then, firing rates were binned with a sliding window (width = 100ms, step = 10ms) and z-scored by subtracting the mean firing rate across time and trials for each cell and dividing by the corresponding standard deviation.

### RNN task definition

We used the Python package *PsychRNN* (Ehrlich et al., 2021) to define tasks, parameterize RNNs, and train supervised RNNs. We used a basic vanilla RNN architecture for all tasks, with its dynamics following the equation:

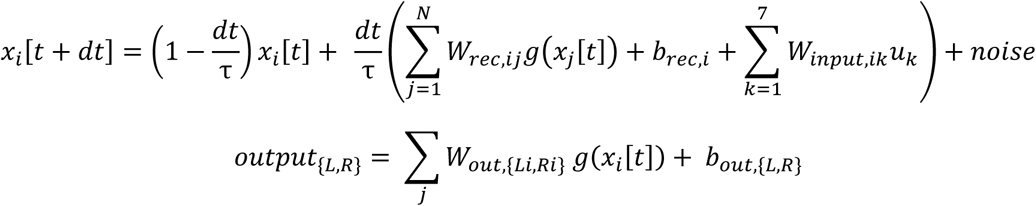

where *g*(*x*) = max(*x*, 0), and the intrinsic time constant *τ* = 100ms, and the simulation timestep *dt* = 10ms. White Gaussian noise was added to both external input *u* (std = 0.01) and synaptic input x (std = 0.1).

To isolate abstract choice formation from spatial action selection, we adapted the task paradigm from Ballesta & Padoa-Schioppa (Ballesta and Padoa-Schioppa, 2019), which involved choice-to-action mappings linking juice types to actions (e.g., Juice A-Left/Juice B-Right). We introduced two additional mappings linking offer order to actions (e.g. Offer 1-Left/Offer 2-Right) to define the “order” task reported in the main text (Fig. 1D). Unlike our animal task in Fig. 1A, the offers were presented at the center of the screen, and both offers were followed by the delay period. No action plan could be made until the mapping cue was shown. We define the presentation of Offer 1 and its subsequent delay as Phase 1, and the presentation of Offer 2 and its subsequent delay as Phase 2. Phase 3 extends from the onset of the choice-to-action mapping cue to the choice response. RNNs trained on Ballesta’s original “juice” task were reported in Figs. 7-8.

Order-task and juice-task RNNs consist of *N* = 50 recurrent units, 7 input channels, and 2 output channels. The output channels model the left (L) and right (R) action (Fig. 1E). The input channels included two juice channels receiving juice quantity normalized by the maximum quantity, four choice-to-action mapping channels (two per task, see the section below), and one fixation channel, which was active until 500 ms after the onset of the choice-to-action mapping cue. Note that two juice types were assumed to be explicitly separated into two channels with independent input weights (Fig. 1E), while the order identities were not externally cued but needed to be inferred through time.

We also modeled the monkey task directly in Fig. 10. There were no choice-to-action mapping channels. Instead, one of two spatial channels was active during each offer presentation, indicating the spatial location of the presentation side and the saccade target.

### RNN training procedure

We trained RNNs with supervised learning algorithm to mimic the behavior choice pattern in monkeys (Fig. 1B). As the relative amount of Juice B, *q_B_*/*q_A_*, increases, choice probability for B rises, yielding a psychometric curve modeled as a logistic function of *q_B_*/*q_A_* (Padoa-Schioppa, 2022):

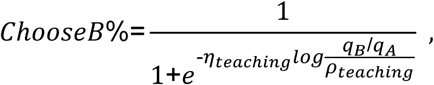

where the indifference point *ρ_teac_*_*hing*_=1.7 and the slope *η_teac_*_*hing*_ = 13 were set as observed from typical experimental sessions. For instance, in a BA trial with (*q_A_*, *q_B_*)=(2,4), the curve yields an 89.2% probability of choosing B.

Using the teaching psychometric curve, we stochastically sampled the “correct” choice for the RNN to learn in each training trial. The juice choice was mapped to the spatial action with the trial-specific choice-to-action mapping in the juice task. For instance, if the correct choice was sampled to be Juice B and the mapping cue is (A-R, B-L) in this trial, then the “correct” action is L. In the order task, this choice was first mapped to the order frame before mapping to the action with the mapping cue. For instance, the “correct” order choice of choosing B in BA trial would be Offer 1, thus action L if the choice-to-mapping cue is (Offer 1-L, Offer 2-R). Note that the additional juice-to-order step only applies to generating the target output for RNN training. For the RNN or monkey subjects themselves, juice-based choices are not necessary in the order task, as order-based choices are not necessary in the juice task.

The trainable parameters are the recurrent weight matrix *W_rec_*, input weight matrix *W_input_*, constant input *b_rec_*, output weight matrix *W_out_*, output bias *b_out_* and initial conditions *x*[0]. In each training epoch, a batch of 112 trials was generated along with their corresponding stochastically generated “correct” outputs. All trainable parameters were updated once every epoch by TensorFlow’s Adam optimizer to minimize the squared error between the target and actual outputs. Parameters were initialized using the default method (random Gaussian weight with radius 1.1). Empirically, we found that training 100000 trials (∼890 epochs) with a default learning rate (0.001) would normally converge. We determined convergence by both the mean squared errors and psychometric curves (described in the following paragraph). We used custom stopping criteria based on the psychometric curve to prevent over-training.

In each training trial, a pair of juice quantities (*q_A_*,*q_B_*) was sampled from the 14 offer pairs used in one typical experiment session, with *q_A_* ∈ [0,4] and *q_B_* ∈ [0,8]. The 14 pairs are [(0, 1), (0, 2), (1, 0), (1, 3), (1, 4), (2, 1), (2, 2), (2, 3), (2, 4), (2, 6), (3, 2), (3, 3), (3, 8), (4, 4)]. Juice quantities were normalized before being fed into RNNs: 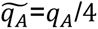 and 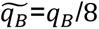, such that 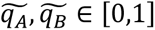. As in the experiments, the presentation order of juices (AB or BA) was randomized across trials, and there was no spatial association during offer presentation, like two offers shown at the center of the screen.

The behaviors of the trained RNNs were measured from the output channel activity from 500-1500ms after mapping cue onset. An offer was defined to be “selected” if its associated output channel had a greater time-averaged activity than the other channel. We calculated the frequency with which juice B was chosen under each offer pair condition and compared it to the expected choice probabilities. A psychometric curve was fit using logistic regression to estimate the RNN’s indifference point *ρ* and slope *η*.

### Training RNN using model-free reinforcement learning (RL)

To test whether the order-based coding observed in supervised RNNs depended on the psychometric-function-derived choice labels, we trained additional RNNs using model-free reinforcement learning, in which networks learned from scalar economic reward contingent on their own actions (Fig. S2; Song et al., 2017). The relative value of Juice A to Juice B was set to 1.7.

These RL-RNNs used an actor-critic architecture. The actor subnetwork served as the decision-making network and received the same inputs as the supervised RNNs, whereas an auxiliary critic network received the actor’s hidden unit activity concatenated with a one-hot representation of the sampled action, and generated a scalar value prediction. Both subnetworks contained 50 recurrent units and shared the same vanilla RNN architecture, activation function, and time constant. Recurrent connectivity was initialized with a spectral radius 1.4, and recurrent synaptic noise was set to 0.15. The recurrent states of both networks were initialized to 0.2 at trial onset, and the critic’s input weights were scaled by a factor of 10 at initialization.

Training used batched Monte Carlo *REINFORCE* with a learned value baseline over 6000 epochs, each comprising 56 parallel trials. Returns were computed as the undiscounted sum of future rewards, and advantages were computed by subtracting the critic’s predicted value from the return, followed by batch standardization. The total loss combined the policy-gradient term, value-function mean-squared error, and an entropy bonus (coefficient 0.03). Actor and critic parameters were jointly optimized using Adam with a learning rate of 0.002 and gradient-norm clipping to 1.0.

### Decoding analysis in RNN and neural data

In each trial, the RNN’s choice could be interpreted in terms of juice identity or order identity. For example, consider a trial in the order task trial with mapping cue = (offer1-L, offer2-R) and BA presentation order. If the RNN selected the Left output, the choice corresponds to juice B, and offer 1. In the experimental task, each offer was characterized by juice type, presentation order, and spatial locations, allowing choices to be labeled under three different reference frames.

To quantify the choice representation in RNN activity, we trained linear discriminators to decode choice labels from population activity snapshots (Fig. 1G-H). Decoding performances are evaluated using 5-fold cross-validation, implemented with *cross_val_score()* and *LinearDiscriminantAnalysis()* function from Python package *Scikit-learn*. To avoid overfitting, we decode the choices from the noisy simulation of sparsely sampled trials, the same batch of 112 trials during RNN training.

In Fig. 1C, the preprocessed experimental data underwent pseudo-population decoding in MATLAB. For each resampling, a pseudo-population was constructed by randomly selecting 20 trials of each label for all cells, splitting into 5 cross-validators with 4 trials each, and building training-test sets through the leave-one-fold-out cross-validation. Linear discriminant classification (*classify* in MATLAB) was then run to achieve the accuracy of label prediction. The resampling was repeated 100 times. Significance of decoding accuracies (DA) at each time bin was tested, in which raw DA (100 resamples) were compared to shuffled DA (100 resamples) at the significance level of p<0.001 by one-sided Wilcoxon rank sum tests.

### RNN single unit analysis

In Figure 2, we examined how RNN units encode the value of Offer 1 in Phase 1 and represent choices in Phase 2. Offer 1 values were defined as *q_B_* for Juice B and *ρq_A_* for Juice A, where *ρ* is each RNN’s behavior indifference point. To determine the tuning of each unit to Offer 1 value, we performed linear regression separately for trials involving each juice type. Time-averaged activities in the last 200ms of Phase 1 were used for linear regression.

**Figure 2.**
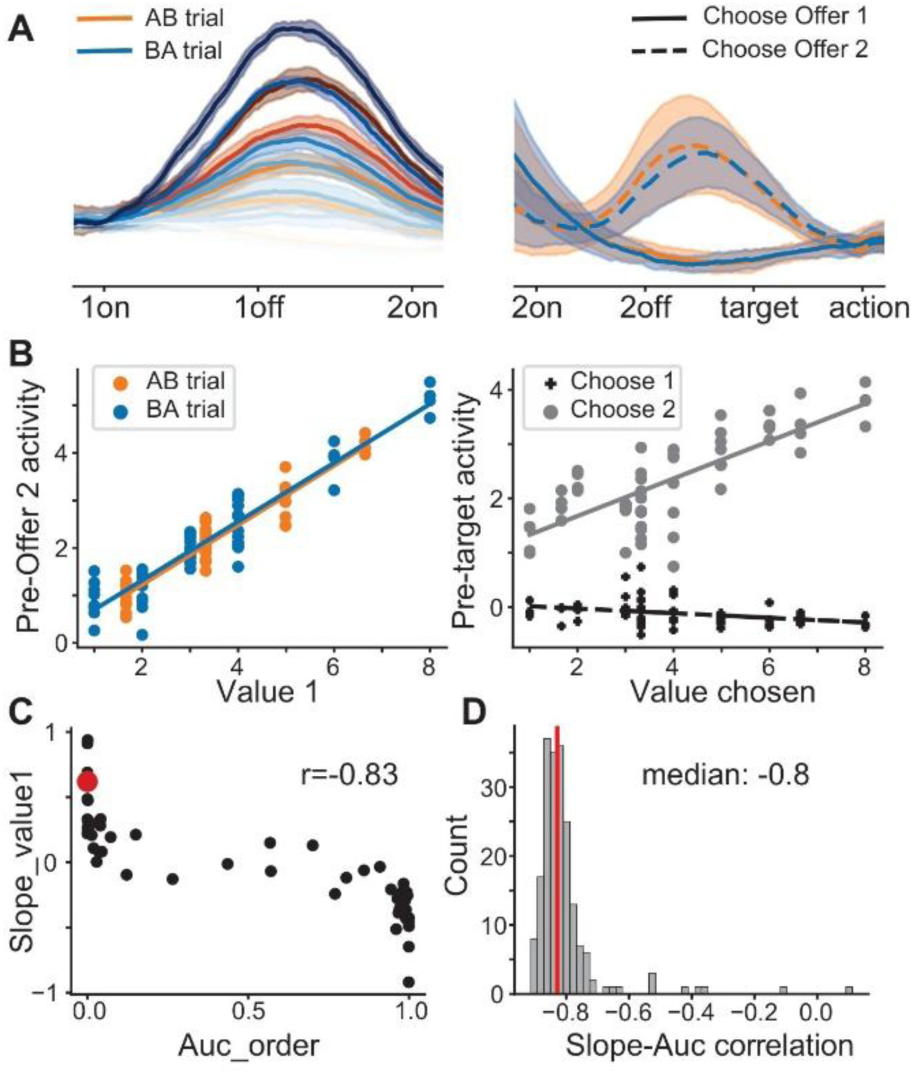
Single units in the order-task RNN. (A) Trial-averaged activity of example units in Phase 1 (Left) and Phase 2 (Right). In Phase 1, activity is grouped by juice types (orange: Juice A; blue: Juice B) and Offer 1 quantity (darker color for larger amounts). In Phase 2, activity is grouped by order choices (solid: choose Offer 1; dashed: choose Offer 2). Shaded areas: ± SD. (B) Tuning to Offer 1 value during the last 200 ms of Phase 1 (Left) and to the chosen value during the last 200 ms of Phase 2 (Right). (C,D) Correlation between encoding and choice preference in the example RNN (C; AUC_order =_ 1/0 corresponds to choose Offer 1/2; the unit shown in A and B is highlighted in red) and across RNN ensembles (D).

To determine choice preference in each reference frame, we conducted Receiver Operating Characteristic (ROC) analysis and computed Area Under the ROC Curves (AUC) for each single unit. We used *roc_auc_score()* function from *scikit-learn*. For the order reference frame, classifiers were defined such that the higher activity was associated with a choice of Offer 1, so an AUC > 0.5 indicates preference for choosing Offer 1, and vice versa. For the juice reference frame, classifiers associated high activity with a choice of Juice A, so units preferring choosing Juice A have AUC>0.5, and vice versa.

### RNN population analysis

To examine the low-dimensional representation and dynamics of trained RNNs, we conducted principal component analysis (PCA) on population activity (Figs. 3,7-9). To capture task-related variability across time and conditions, RNN activity from Phases 1 and 2 was concatenated across trials to form a data matrix *X*. We used the PCA() function from *scikit-learn* to estimate the principal components (PCs). We projected population activity onto the first two PCs for visualization. To visualize recurrent dynamics, we computed the derivatives of the input-free RNN dynamics at a grid of points in the PC1-PC2 plane (with all other PC dimensions fixed to zero) and plotted the resulting vector field.

**Figure 3.**
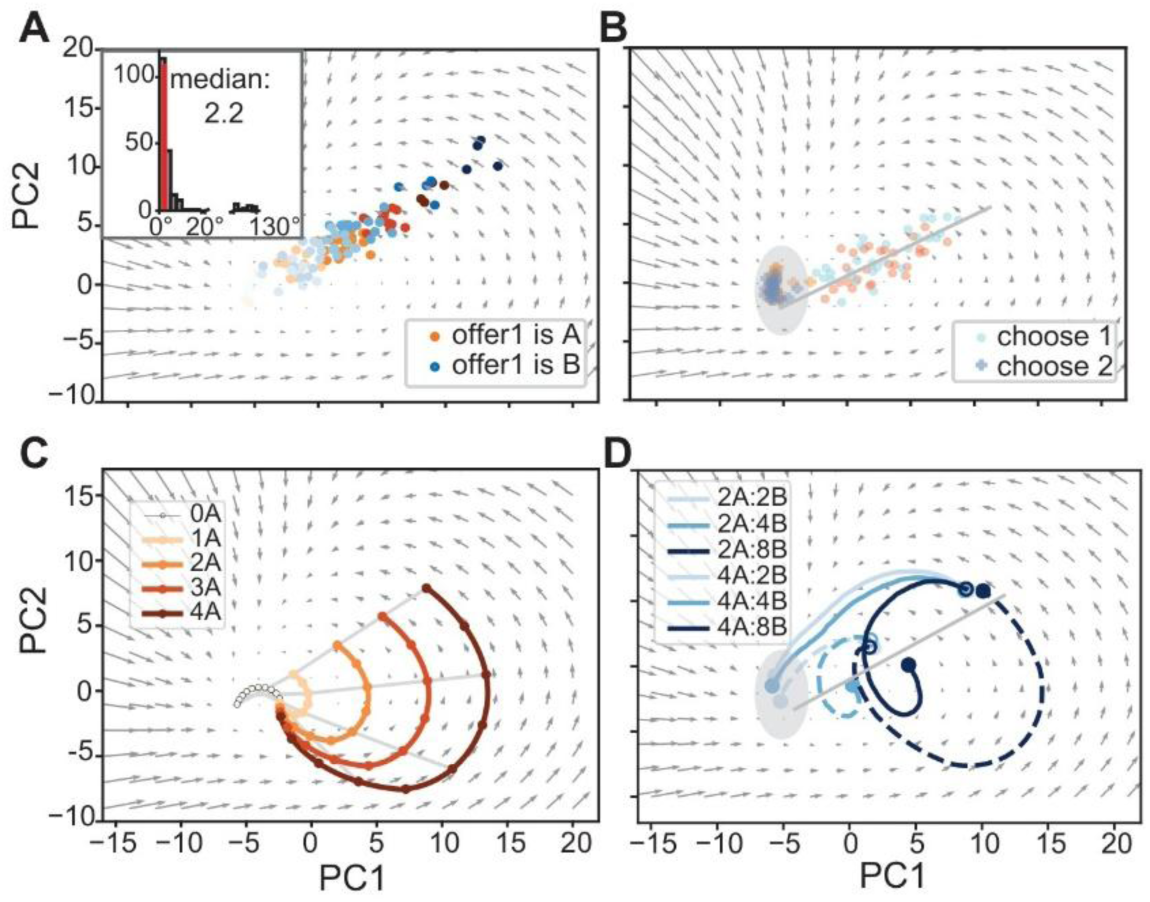
Population analysis. (A,B) Example RNN activity at the end of Phase 1 (A) and Phase 2 (B), projected onto the first two principal components (PCs). Color indicates juice type (orange: Juice A; blue: Juice B), with darker shades representing larger Offer 1 quantities. Gray arrows show vector fields of RNN dynamics without input. At the end of Phase 2, states corresponding to Offer 1 choices cluster within the gray ellipse, whereas those corresponding to Offer 2 choices are distributed along the gray line (B). (*Inset*) Distribution of angles between encoding axes for the two juices across RNNs. (C) Trial-averaged Phase 1 activity. For simplicity, only AB-trial conditions with juice A in Phase 1 were shown. Gray lines connect points 100ms apart during the inter-offer period. (D) Trial-averaged Phase 2 activity, with traces starting from open circles and ending at filled circles. A subset of AB trials from (C) is shown to illustrate how choices diverge depending on the initial condition set by Offer 1 memory (open circles) and Offer 2 value (*q_A_* =2: dashed; *q_A_* = 4: solid; *q_B_* =2,4,8: light, medium, and dark).

To assess the geometric relationship between juice representations in Phase 1 (Fig. 3A), we measured the angle between two juice coding axes. These axes were defined using the regression coefficients: for each juice type, we extracted the linear regression slopes with respect to the Offer 1 value across 50 units, forming two 50-dimensional vectors. The angle between them was measured as the inverse cosine of their normalized inner product.

### Latent circuit inference and analysis

To gain mechanistic insight into RNN dynamics (Figs. 4 and 7D), we adapted the method from (Langdon and Engel, 2025) to infer a low-dimensional circuit embedded in the high-dimensional system. Unlike Langdon and Engel, though, we didn’t restrict each node in the latent circuit to selectively encode any input channel. As we were more interested in abstract choice formation rather than the choice-to-action formation, we restricted the analysis to Phases 1 and 2 and dropped the output.

**Figure 4.**
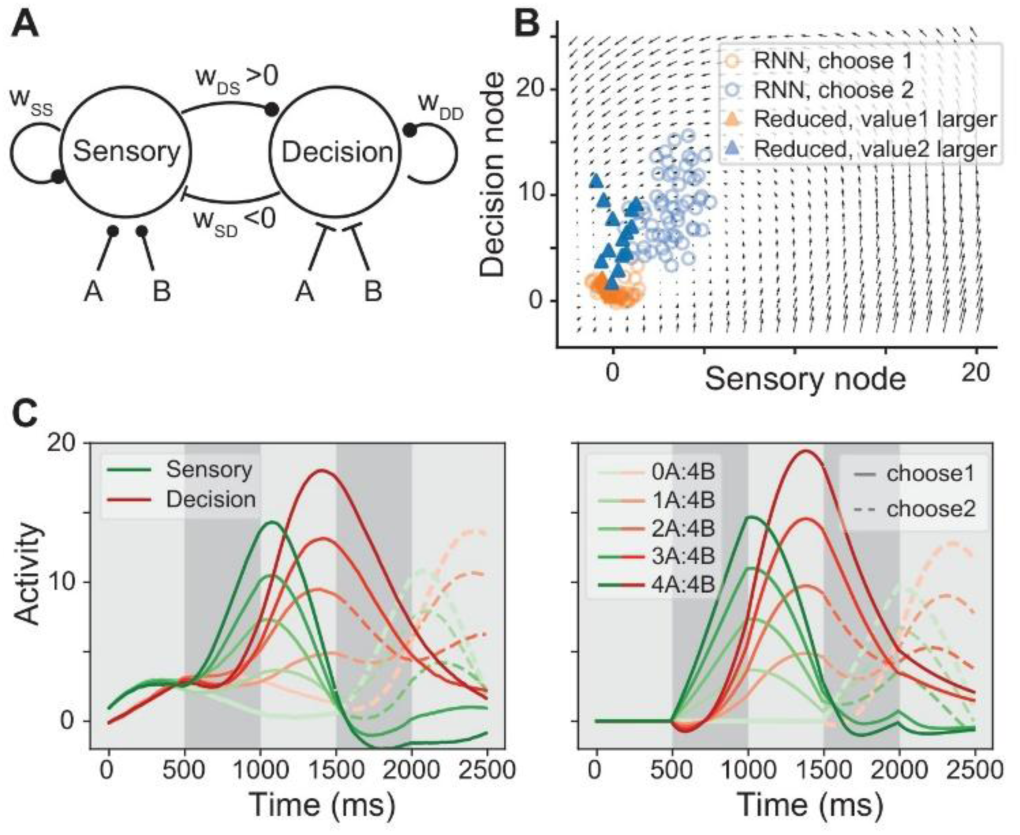
Latent circuit inference. (A) Reduced circuit diagram with inferred weights of the example network. Recurrent: 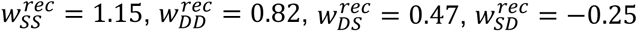. Input; 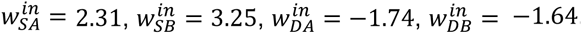 (B) Consistency between the RNN and its reduced circuit. RNN activity was projected onto the two nodes using the inferred embedding matrix *Q* (Methods); vector fields were obtained from the reduced circuit’s dynamics. (C) Trial-averaged activity projected onto the two nodes for AB trials. Left: RNN projection *Q^T^x*(*t*). Right: activity of the reduced circuit *y*(*t*). Trials with fixed *q_B_* = 4 are shown; color darkness indicates varying *q_A_*. Solid and dashed lines represent trials with higher choice probability for Offer 1 and Offer 2, respectively.

Here, we approximate RNN activity 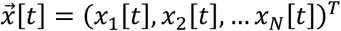 (time *t* within Phases 1 and 2) using a two-dimensional dynamical system 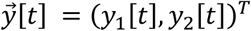 and an orthonormal embedding matrix Q ∈ ℝ*^N^*^×2^ such that

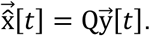

The model was trained to minimize the sum of mean squared error ∑*_i_*_,*t*_(*x_i_*[*t*] − *x*^*_ι_*[*t*])^2^.

The dynamics of the reduced circuit are governed by

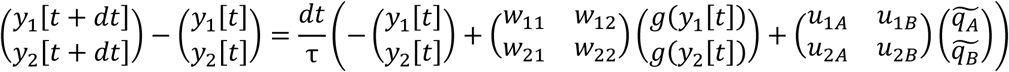

where *w_ij_* is the recurrent weight matrix, *u_ij_* is the input weight matrix, *q̃_ι_* is the normalized input, and the nonlinearity function *g*(*y*) = max(*y*, 0). Parameters *Q*, *w_ij_*, and *u_ik_* (*i*, *j* ∈ 1,2, *k* ∈ *A*, *B*) were optimized using custom Python code with PyTorch with Adam optimizer (learning rate = 0.001, weight decay = 0.001). Optimization was stopped when the loss had not improved by more than 0.001 over 25 epochs. For each RNN, we repeatedly initialized parameters and optimized them until the reconstruction loss fell below the variability of RNN activity.

We examined the effect of perturbing network or task parameters on the choice order bias (Fig. S5) (Padoa-Schioppa, 2022). We fit RNN choices separately for AB and BA trials to obtain indifference points *ρ_AB_* and *ρ_BA_*. The order bias *ε* is defined as:

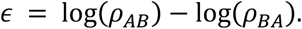

A bias towards Offer 1 yields a lower *ρ_AB_* and higher *ρ_BA_*, resulting in a negative *ε*.

### Correlation between regression coefficients in RNN and neural data

In Figure 5, we calculated the Pearson correlation coefficient between regression weights. Time-averaged activities in the entire offer presentation period in the neurophysiological data and in the last 200ms in RNN were used for linear regression. As noted by (Ballesta and Padoa-Schioppa, 2019), spurious value tuning can be an artifact arising from unequal value ranges across juices and correlation between offer values *v*_1_ and *v*_2_. To address this issue, we used a restricted set of offer pairs with no correlation between *v*_1_ and *v*_2_. Specifically, we included two trial types, either *q_A_*= 2 with varying *q_B_*, or *q_B_*= 4 with varying *q_A_*, noting that 2A is approximately equivalent in value to 4B. This design yields a cross-shaped distribution of offers without correlations. For monkey F, in sessions where the indifference points *ρ* deviated substantially from 2, we instead fixed *q_B_* at the value most comparable to 2A, rather than at 4B. For the neurophysiological data, we further restricted the analysis to cells that were task-selective in at least one consecutive 500-ms task epoch, based on an ANOVA test (p<0.01), resulting in 285 and 152 OFC neurons in monkey N and F, respectively.

**Figure 5.**
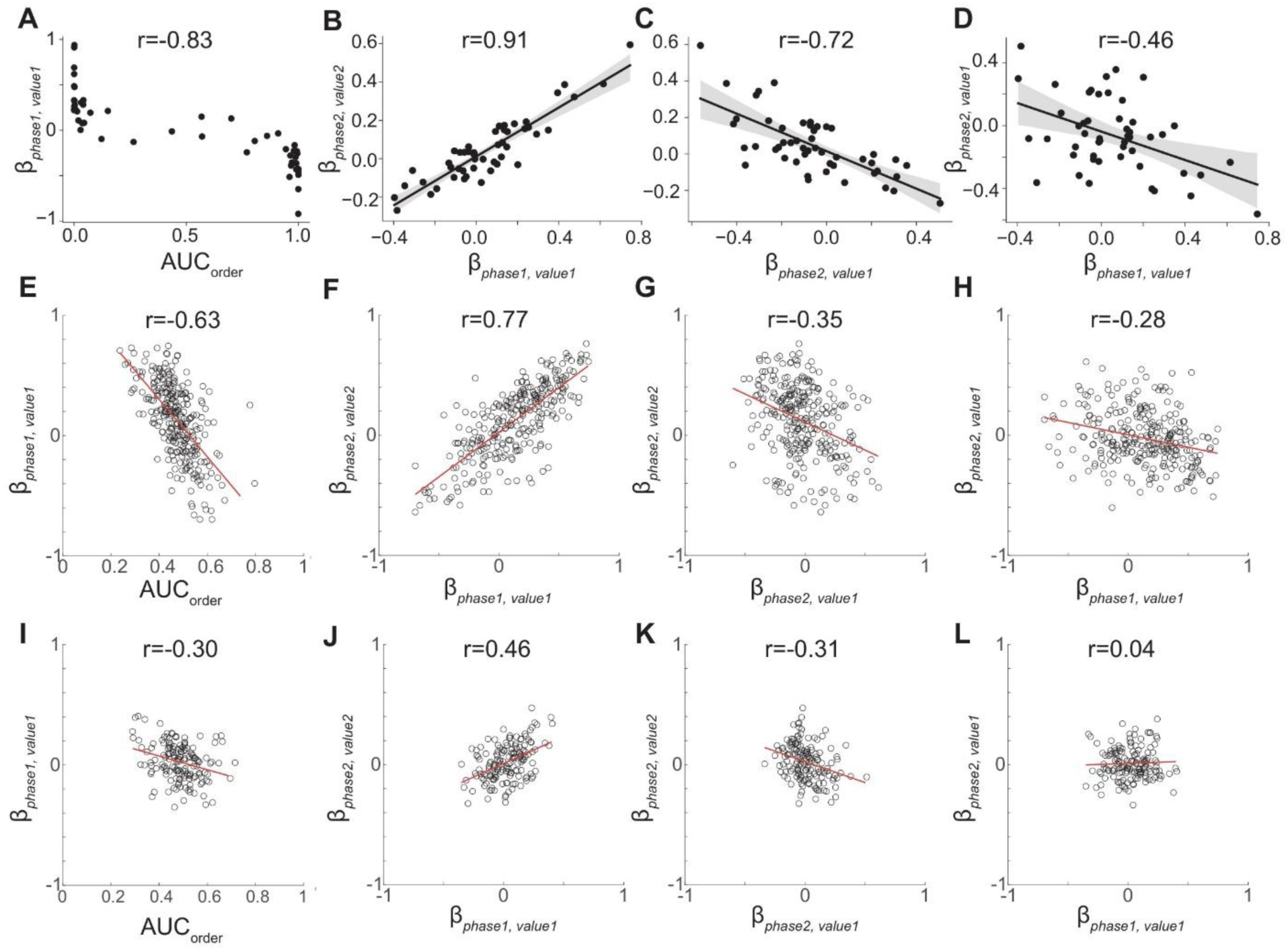
Neural data in monkey experiment consistent with order-task RNN. (A-D) Order-task RNNs. A: correlation between Offer 1 tuning in Phase 1 and order choice preference in Phase 2; B: Offer 1 tuning in Phase 1 vs. Offer 2 tuning in Phase 2; C: Offer 1 and Offer 2 in Phase 2; D: Offer 1 tuning in Phase 1 and Phase 2 (*N* = 50). (E-L) Same layout as in (A-D) but from monkey data (E-H: Monkey N, *N*=285; I-L: monkey F, *N*=152). All Pearson correlation P < 0.001 except for **L** with *P* = 0.6185.

### Cosine similarity between offer-value coding axes

In Figs. 6 and 10, we analyzed epoch-averaged activity during first-offer presentation (FO), early and late inter-offer delays (ED and LD), second-offer presentation (SO), and the second delay (SD). The experimental inter-offer delay was twice as long as that in the RNN task, and the experiment had no second delay. During Phase 1, each unit’s activity was regressed on Offer 1 value; during Phase 2, both offer values were included in the same regression. Vectors of the corresponding regression coefficients across units defined the offer-value encoding axes. Subscripts indicate the encoded offer value; FO, ED, and LD refer to Offer 1 encoding.

**Figure 6.**
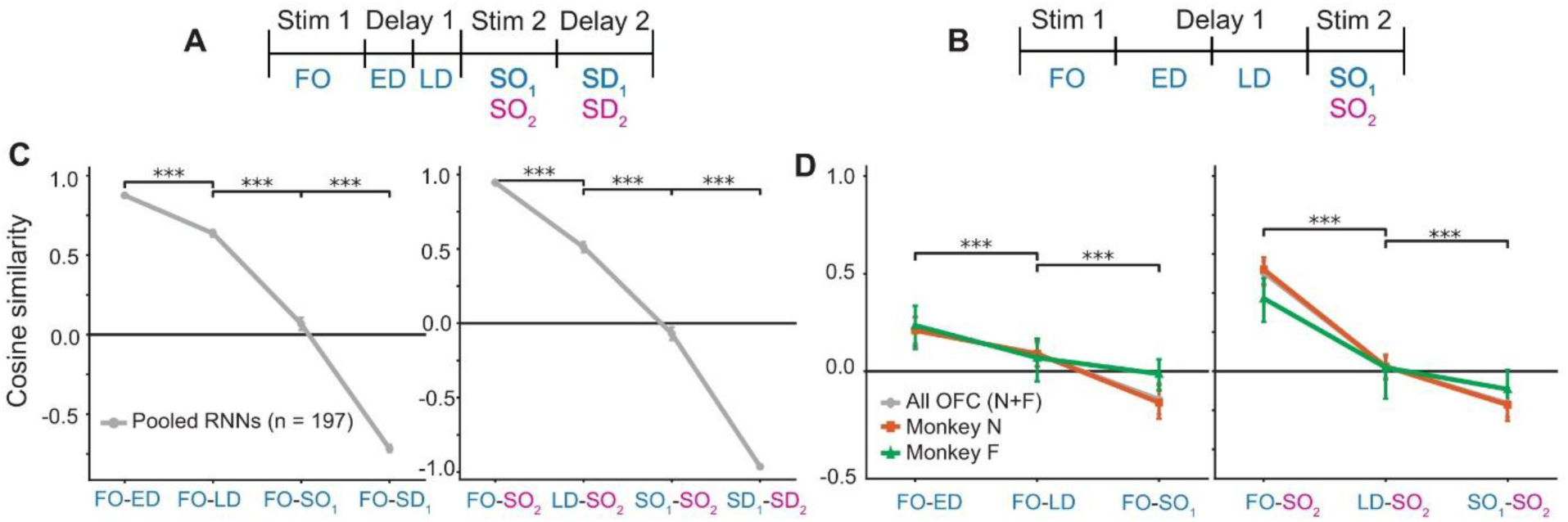
Cosine similarity of offer-value encoding directions across task epochs. (A,B) Task epochs in RNNs (A) and the experiment (B). FO, first-offer presentation; ED/LD, early/late inter-offer delay; SO, second-offer presentation; SD, second delay. Subscripts indicate the encoded offer value in Phase 2, and FO, ED, and LD refer to Offer 1 encoding. The experimental inter-offer delay was twice as long as that in the RNN task, and the experiment had no second delay. (C,D) Cosine similarities between Offer 1 encoding directions across epochs (left) and between Offer 1 and Offer 2 encoding directions (right), for RNN (C; n=197) and OFC pseudo-populations (D; pooled, gray, n = 1226 neurons; monkey N, orange, n = 824; monkey F, green, n = 402). Points indicate mean similarities; error bars indicate 95% bootstrap confidence intervals obtained by network bootstrap for RNNs and session-block bootstrap for OFC. Brackets indicate comparisons between adjacent plotted similarities (OFC: pooled data only); ***p < 0.001, two-sided paired bootstrap tests.

**Figure 7.**
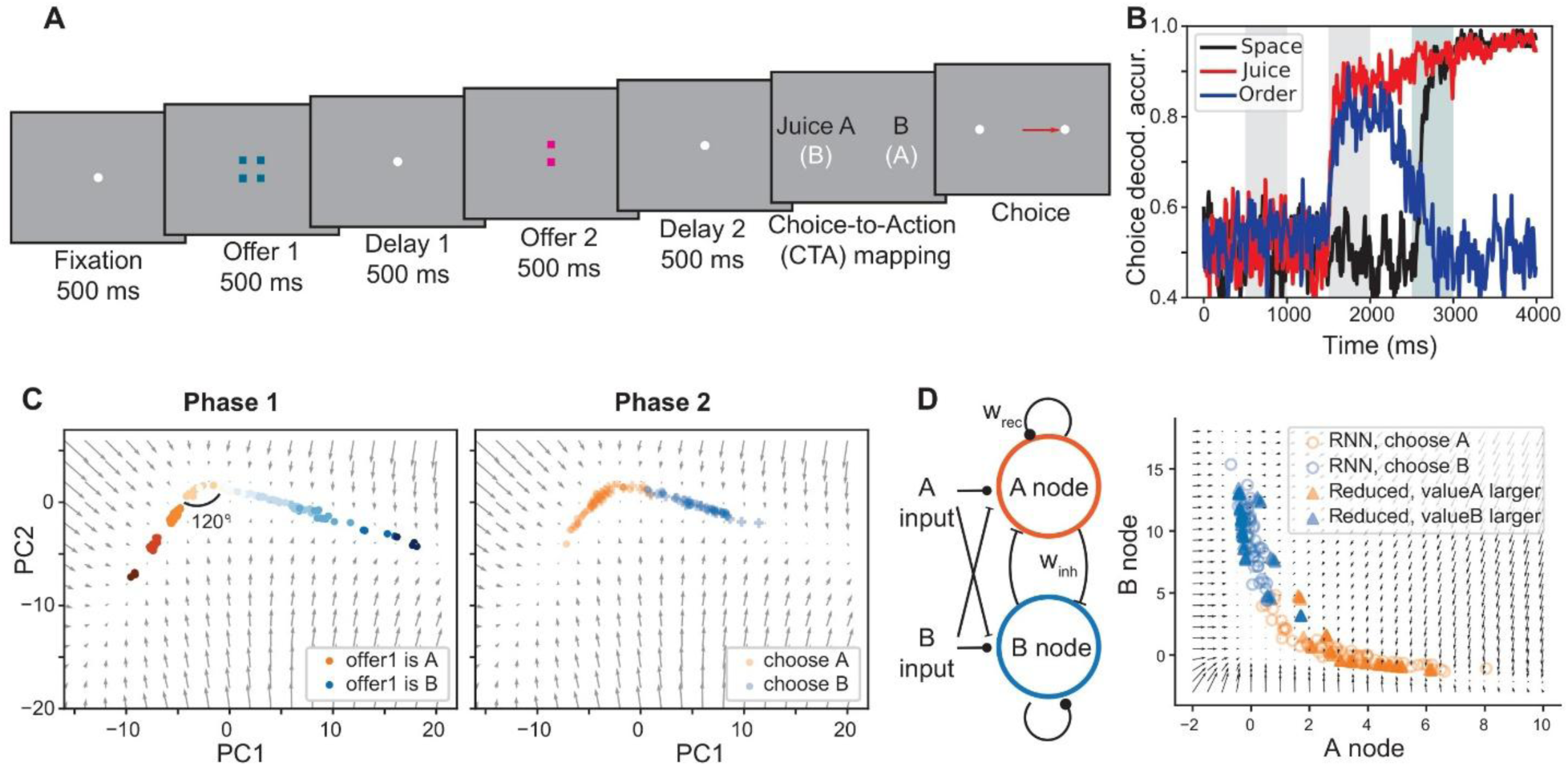
RNNs for juice-based choice signals and the emergence of attractor dynamics. (A) Task design for RNN. (B) Choice decoding analysis confirmed juice-based choice signals. (C) Example snapshots of juice-task RNN activity at the end of Phase 1 (Left) and Phase 2 (Right), projected onto the top two principal components (PCs). Color indicates juice type (orange: Juice A, blue: Juice B) with darker shades representing larger Offer 1 quantities. Background vector fields (gray arrows) show RNN dynamics in the absence of input. (D) Reduced circuit diagram of the juice task, showing competitive nodes responsible for encoding and choosing different juices. Inferred weights of the example network – recurrent: 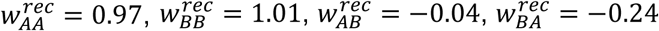; input weights: 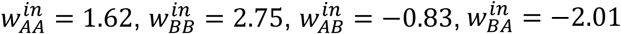. Consistency of choice representations and vector fields between the juice-task RNN and its reduced circuit: RNN activity projected onto the A and B nodes using the inferred embedding matrix Q (Methods), with vector fields obtained from the reduced circuit’s dynamics.

**Figure 8.**
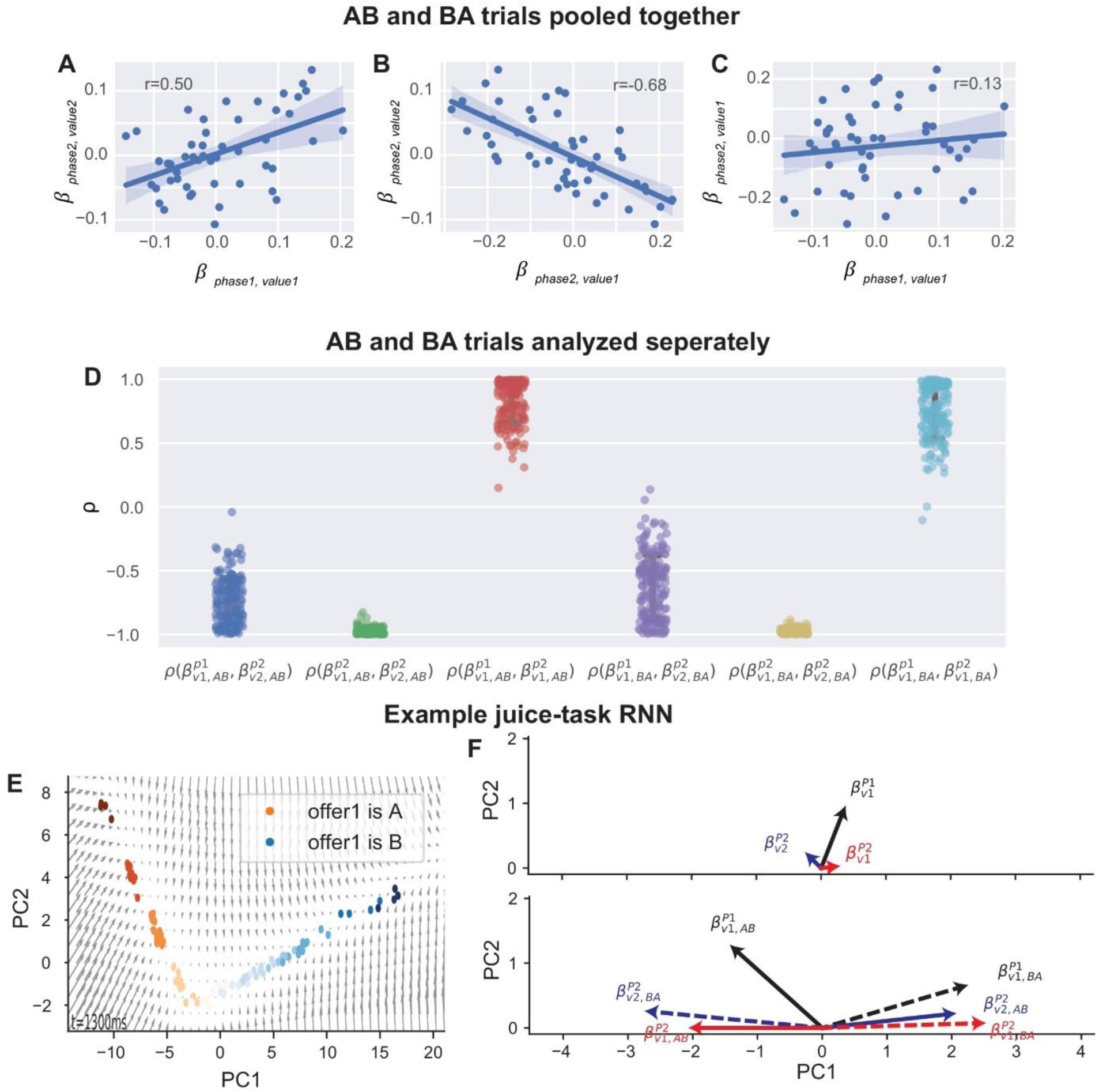
Correlation patterns in the juice task when AB and BA trials were pooled together (A-C) or analyzed separately (D-F). (A-D) Correlation coefficients of regression weights for Offer 1 and Offer 2 values across all trials (A-C) and in AB (D, left) and BA trials (D, right). In all cases, 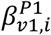 and 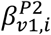 are positively correlated, reflecting stable value representation (C,D), and 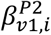 and 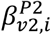 are negatively correlated, because of mutual inhibition (B,D). Notably, the correlation pattern between 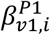 and 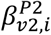 differs when AB and BA trials are combined (A) versus analyzed separately (D). (E,F) Example RNN illustrating variable correlation patterns when AB and BA trials were pooled. (E) Snapshots of value representation at the end of Phase 1, showing similar patterns across RNNs. (F) Regression coefficient vector 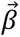 projected onto the PC planes for Offer 1 in Phase 1 (black), Offer 1 in Phase 2 (red), Offer 2 in Phase 2 (blue), averaged over AB and BA trials (top; Methods) or shown separately (AB: solid, BA: dashed). The vectors in the top panel are the average of those in the lower panels. In Phase 1, 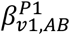 and 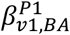 align with the A and B axes. In Phase 2, activity spreads across both axes (see Fig. S4B), tilting Offer 1 and Offer 2 value representations away from these axes and making them nearly antiparallel in AB and BA trials. This reduces regression coefficient strengths (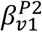 and 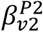) in the top panels and creates large fluctuations in correlation coefficients.

For each RNN, regressions were fitted to training-fold trials using three stratified folds, separately for AB and BA trial configurations. Coefficient vectors were pooled across folds and configurations for each axis, and axis similarity was calculated as the cosine between the pooled vectors. Similarities were then averaged across networks. For OFC neurons, epoch-averaged firing rates were baseline-subtracted using the pre–Offer 1 window and z-scored across trials for each neuron. Encoding axes were estimated using the same regression models as for RNNs. To reduce shared-trial noise effects, trials were divided into odd and even subsets, and each axis was refitted separately in each half. Similarities were averaged across split-half comparisons. OFC pseudo-populations were analyzed separately for each monkey and for the pooled population. Population 95% CIs and paired differences between neighboring pairs (pooled monkeys only) were obtained by session-block bootstrap: sessions were resampled with replacement, and cells of a session moved together. p-values were two-sided percentile-based against the null of equal cosines.

### Code accessibility

All RNN training and analysis codes are available on GitHub (https://github.com/jtg374/EconomicDecisionMakingReferenceFrame).

## Results

### Order-based choice in sequential-offer economic decision-making

To investigate the mechanism of order-based choice representation in economic decision-making, we considered a juice task where the monkey needs to choose between different types and quantities of juices (Fig. 1A). The two offers were presented sequentially on different sides of the screen, while the order of presentation and spatial locations were randomized.

Behavioral choices can be summarized by a psychometric curve relating the probability of choosing Juice B to the ratio between the offered quantities of Juices A and B, with A being preferred (Fig. 1B). As the relative amount of juice B increases, the probability of choosing B rises, and the indifference point – where the animal chooses between A and B with equal probability – occurs when the amount of B exceeds that of A. Behavior choices in this task are largely independent of presentation order or the spatial side of the offers.

Because each offer contains multiple attributes – including juice type, quantity, presentation order, and spatial sides – choice variables can be represented in several reference frames: juice identity, order, and space (Padoa-Schioppa and Conen, 2017; Cai and Padoa-Schioppa, 2019). Notably, decoding analyses of OFC activity revealed that after the second offer presentation, order-based choice emerged first, followed by a transformation into spatial choice signals that guide action, whereas juice-based representations are relatively weaker (Fig. 1C). This differs from typical juice tasks where juice-based choice dominated (Padoa-Schioppa, 2011; Padoa-Schioppa and Conen, 2017). Notably, previous work has highlighted substantial overlap between value representations and order-choice signals in the OFC (McGinty and Lupkin, 2023). Here, we examine this relationship in the present sequential juice-choice task and use the model to provide a mechanistic account.

### Task-optimized RNN with order-dependent behavior reporting

To identify the circuit mechanism of working memory and decision-making underlying order-based choices, we trained task-optimized RNNs (Barak, 2017; Ehrlich et al., 2021). To isolate abstract order choice formation from spatial action selection, we modified the task so that the spatial information was unavailable during offer presentations. Instead, after the presentations of two offers, the network received a choice-to-action mapping cue specifying how the order of the chosen offer should be translated into a spatial action (e.g. first/second with left/right or vice versa; Fig. 1D). This manipulation allowed us to examine order-based computations separately from their translation into spatial actions, making the underlying dynamics and latent circuits more interpretable. The same framework also provided a controlled comparison with identity-based choice: in juice-task RNNs, the mapping cue linked spatial actions to juice identity rather than presentation order, allowing us to examine how circuit mechanisms differ across choice reference frames (see “Comparison to identity-based choice” in Results and Discussion).

The task therefore consisted of three phases: Phase 1 presents the first offer (Offer 1) followed by a delay (inter-offer period), Phase 2 presents the second offer (Offer 2) followed by another delay (wait period), and Phase 3 requires a left or right action with the mapping cue. The RNN received five input channels and produced two output channels (Fig. 1E). Two input channels encoded the quantities of Juices A and B during the offer presentations. Two additional channels specified the choice-to-action mapping during Phase 3, and a fixation channel remained active until 500 ms after the mapping cue onset. The network’s two output channels reported either left or right choices in Phase 3.

We trained RNNs to reproduce the behavior choice patterns observed in monkeys (Fig. 1F). The target psychometric curve was defined with the indifference point of 1.7. For each juice ratio, the RNN’s target output was randomly sampled according to the behavior choice probability, introducing stochasticity in the teaching signal. The trained RNN reproduced psychometric curves with indifference points close to the target value. For each trained network, we inferred the subjective value of each juice from the fitted psychometric curve. In the example RNN shown in Fig. 1F, the estimated indifference point is 1.61, indicating that Juice A has approximately 1.61 times the value of Juice B for the same quantity. Offer values were defined in Juice B units using the indifference point.

The trained RNN exhibited dominant order-based choice signals beginning at the onset of Offer 2, followed by spatial choice representations emerging after the mapping cue in Phase 3 (Fig. 1G). In contrast, juice-based choice signals were generally weak, although a few trained RNNs showed coexistence of juice-based and order-based representations (Fig. 1H; see also Fig. 9). These results confirmed that the choice reference frame in the RNN can be shaped by the output-contingent mapping, and we refer to RNNs trained to show order-based choice signals as order-task RNNs.

**Figure 9.**
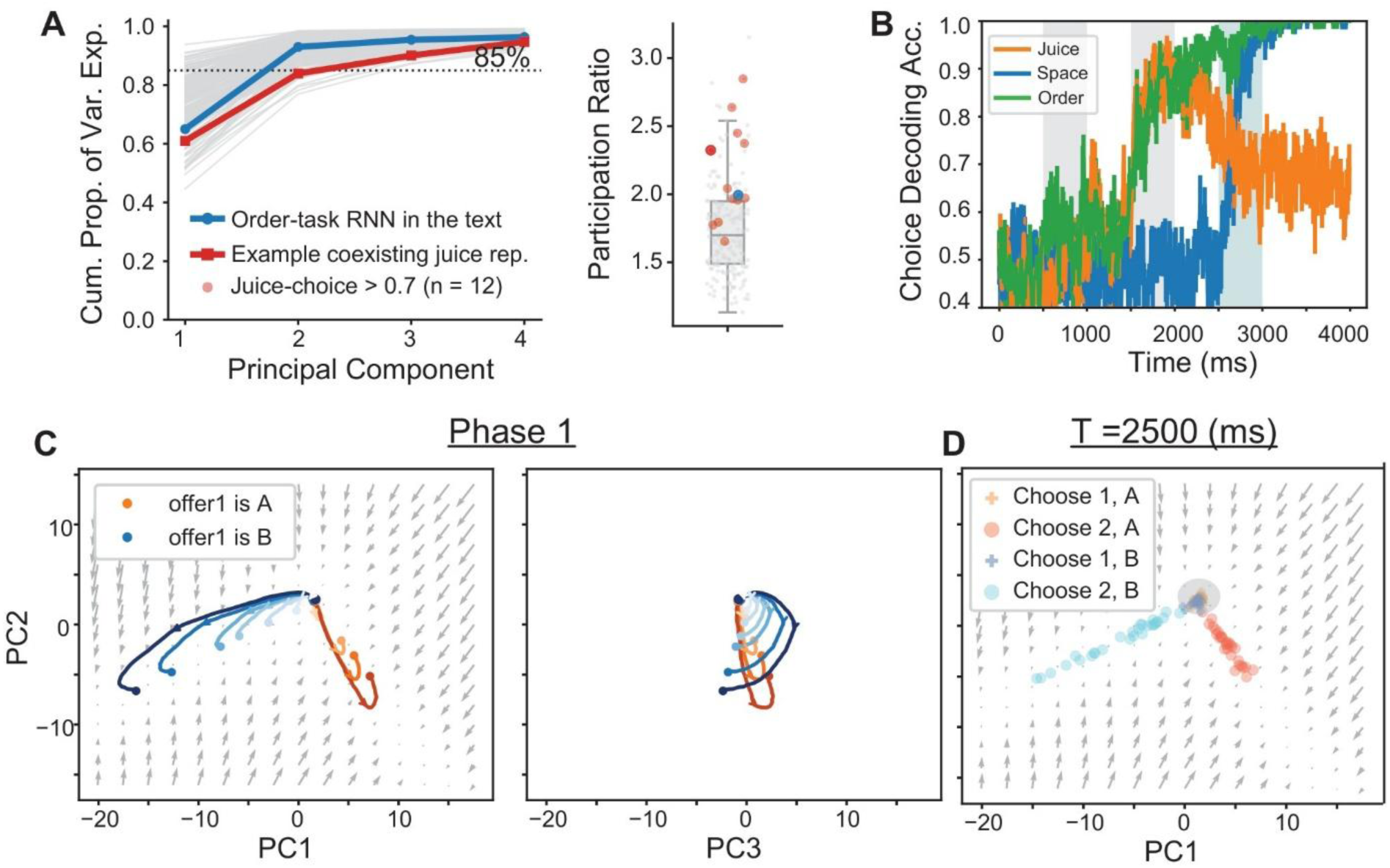
Coexisting order-based and juice-based choice representations in an order-task RNN. (A) Cumulative variance explained by principal components (left) and participation ratios (right) across order-task RNNs. The example in Figs. 2-5 is highlighted in blue, and those showing high juice-choice decoding are shown in red. (B) Choice-decoding accuracy for the example order-task RNN highlighted in red, which exhibited high choice-decoding signals in both juice- and order-based frames. These networks tend to have transient juice-frame choice representation and remain decodable above chance through the task. (C) Phase 1 trajectories projected onto the PC1–PC2 (left) and PC3–PC2 (right) planes, showing convergence toward juice-specific subspaces (left) and prolonged transient dynamics (right) during the inter-offer delay. The vector field is omitted from the PC3–PC2 projection because the projected dynamics largely depend on the position along PC1. (D) Choice representations at the end of Phase 2 (t=2500ms). As in the example order-task RNN, Offer 1 choices clustered near the branch origins (gray ellipse), whereas Offer 2 choices extended along the branches. Juice-based choice representations approximately occupied the juice-specific subspaces identified during Phase 1.

**Figure 10.**
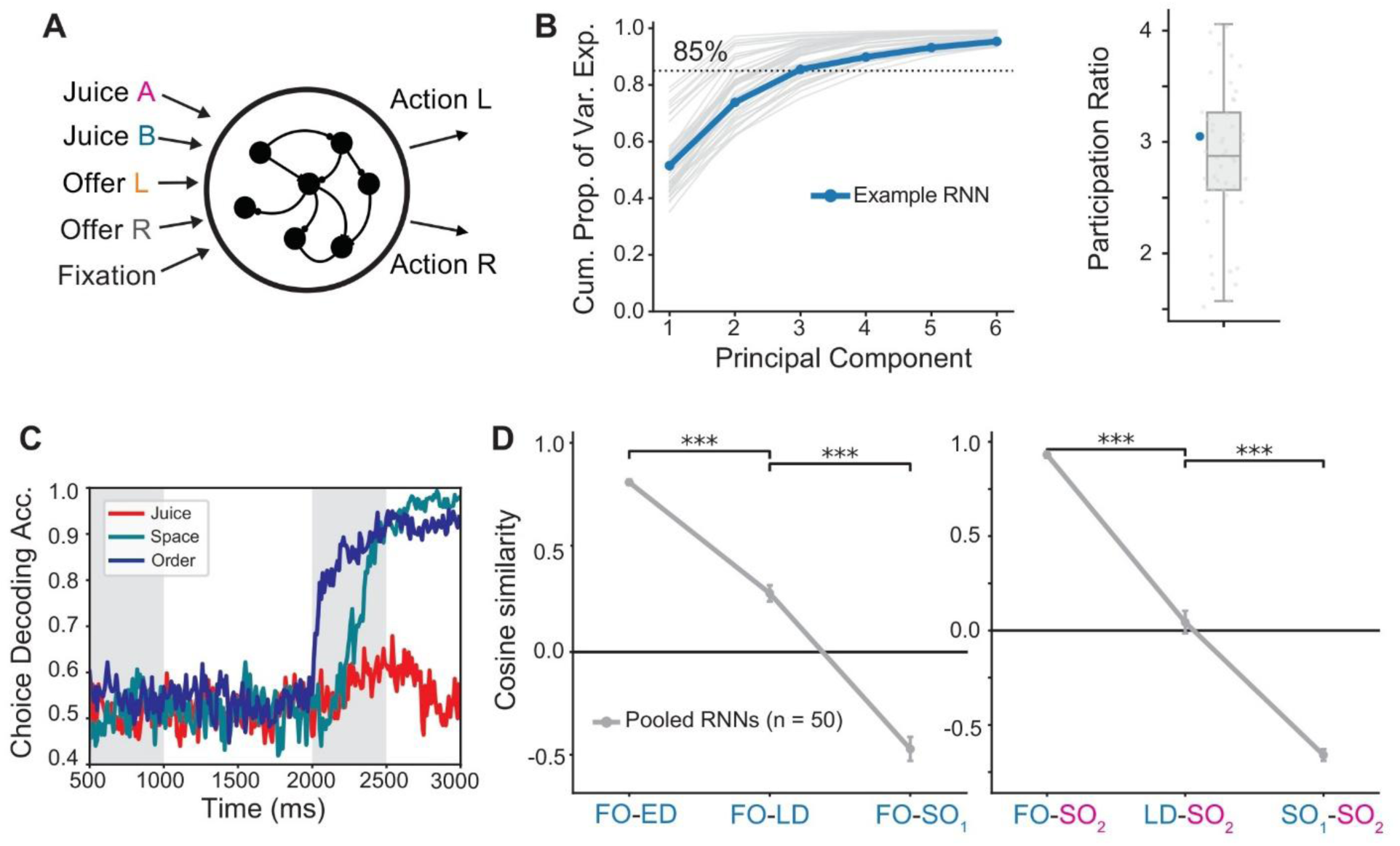
RNNs trained on the monkey task exhibit similar representational signatures of order-task RNNs. (A) RNN architecture. During each offer presentation, one of two spatial channels is active, indicating the offer’s location. (B) Cumulative variance explained by principal components (left) and participation ratios (right) across monkey-task RNNs. Most networks had participation ratios greater than 2. (C) Choice-decoding accuracy in the example monkey-task RNN. Order-based choice signals emerged rapidly after Offer2 onset, preceding spatial-based choice signals, as in the monkey data. (D) Mean cosine similarities between Offer 1 encoding directions across epochs (left) and between Offer 1 and Offer 2 encoding directions (right) across 50 RNNs, showing transformations qualitatively similar to those in OFC and order-task RNNs. Encoding directions are labeled using the same notation as in Fig. 6. Brackets indicate comparisons between adjacent plotted similarities; ***p < 0.001, two-sided paired network-bootstrap tests, 1,000 resamples.

### Juice-agnostic encoding and reversed choice preference

To investigate the mechanisms underlying order-based choices, we first analyzed single-unit activity of order-task RNNs. Specifically, we examined how tuning changes before and after the presentation of Offer 2, which reveals how Offer 1 is encoded and maintained and how it is compared with Offer 2.

In most RNNs, unit activity did not segregate by juice types despite separate input channels for each juice (Fig. 2). During Phase 1, nearly all offer-responsive units exhibited the same tuning sign for both juices. In the example unit, the value tuning for Juices A and B even showed a similar slope (Fig. 2A,B, left). Phase 2 activity was also indistinguishable for choosing A versus B (Fig. 2A,B, right). These findings align with the decoding analysis showing that juice identity cannot be reliably decoded in the order task.

Instead, order signals emerged during Phase 2, particularly as a negative correlation with the value tuning observed during encoding (Fig. 2C,D). For instance, units positively tuned to Offer 1’s value tended to prefer choosing Offer 2 with higher overall firing rates and positive tuning (Fig. 2A,B). To quantify order-based choice preference at the end of the wait period, we used an area under the receiver operating characteristic curve (AUC) score. As illustrated by the example unit, order-based choice preference was negatively correlated with the tuning slope for Offer 1 value across units within a trained RNN (Fig. 2C) and across an RNN ensemble trained on the same order tasks but with different initialization (Fig. 2D).

Note that the negative tuning–choice correlation (Fig. 2C,D) does not by itself establish choice-related activity beyond offer-value encoding. For example, a neuron positively tuned to Offer 1 value in Phase 1 tends also to show positive tuning for Offer 2 value during Phase 2 (Fig. 2A,B). Because higher Offer 2 values increase the probability of choosing Offer 2, this neuron may exhibit higher Phase 2 activity on trials in which Offer 2 is chosen. Since larger AUC_order v_alues indicate an Offer 1 preference, overlapping current-offer value representations across phases can therefore generate a negative correlation without a separate choice-specific component. We next use RNNs to examine the role of these overlapping representations in the mechanism of order-based choice.

### Dynamic and overlapping population activity

We next examined the relationship between value encoding/maintenance and decision-making processes at the population level. Specifically, we applied principal component analysis (PCA) to Phases 1 and 2 to capture most of the activity variability arising from temporal dynamics and differences across conditions, such as trial type and offer values. Across both phases, RNN activity was largely restricted to a two-dimensional subspace. In example order-task RNNs, the first two principal components accounted for 95.7% of the total variance.

Consistent with the single-unit activity, population activity in the example order-task RNN showed overlapping value representations for the two juices during the inter-offer period (Fig. 3A; Video 1). This overlap is reflected in the near-zero angle between the axes encoding the two juices’ values across RNNs (Fig. 3A, inset). Choice representations were also aligned across juices and orders, forming a unified “choice axis” (Fig. 3B; Video 2). Yet, the specific order remained decodable from the relative position along the choice axis. Notably, trajectories corresponding to choosing Offer 2 spread broadly along the axis, while Offer 1 choices were concentrated near its tip, consistent with higher firing rates observed for Offer 2 choices in single-unit analysis.

The overlap in value encoding and choice representations across different juices or orders suggests that the dynamics of order-task RNNs are not governed by attractor mechanisms that support stable working memory and choice representation for different juice types. Instead, value representations evolve dynamically. Following the Offer 1 presentation, the activity grows proportionally to its value along what we term the encoding axis. During the inter-offer period, the activity rotates while preserving value information in the radius of rotation (Fig. 3C). Near the onset of the Offer 2 presentation, the representations of Offer 1 value were transformed into a direction nearly orthogonal to the original encoding axis.

With the onset of Offer 2, trajectories diverge according to its value relative to the remembered value of Offer 1 (Fig. 3D; Video 2). When Offer 2 is relatively less valuable (2A:2B, 4A:2B, and 4A:4B in AB trials), trajectories decays back to the rotation center. In contrast, when Offer 2 is relatively more valuable (2A:4B, 2A:8B, and 4A:8B), its input drives renewed activity along encoding axis, followed by another rotation during the second delay. As a result, the trajectories associated with choosing Offer 1 stay near the rotation center, while Offer 2 choices extend along the choice axis, allowing the order of the chosen offer to be decoded from the distance to the rotation center.

To link the inferred dynamics to behavior, we tested whether trial-to-trial differences in the RNN population state immediately before Offer 2 predicted subsequent choice and whether perturbing this state altered the decision (Fig. S1). Pre-Offer-2 activity was predominantly one-dimensional, and its projection onto the choice-related axis distinguished subsequent choices with an AUC significantly above chance. Perturbing this state along either choice axis or the Offer 1 value-encoding axis (PC1) produced similar effects on subsequent choice. Together, these results demonstrate that the population state reflecting Offer 1 value and internal noise during Phase 1 can influence the decision made in Phase 2.

We further examined whether dynamic features of order-based choices depended on the indifference point or learning objectives. As an alternative to training with psychometric-function-derived choice labels (Fig. 1), we trained additional order-task RNNs using model-free reinforcement learning, with scalar economic rewards contingent on their own choices (Fig. S2). We also changed the instructed indifference point and continued training a previously trained supervised RNN (Fig. S3). In both cases, the principal encoding relationships of Offer 1 and 2 encoding directions remained qualitatively unchanged, supporting the robustness of the coding geometry and dynamics underlying order-based decisions in the model.

### Circuit mechanism underlying order-based choice

To gain mechanistic insight into value encoding and comparison in order-task RNNs, we inferred a low-dimensional connectivity structure (Langdon and Engel, 2025). Using activity from Phases 1 and 2, we derived a reduced circuit consisting of two nodes, yielding matrices that embed latent-circuit activity to RNN activity and capture the strengths of input, output, and recurrent connections (Methods).

The resulting circuit structure was asymmetric in both feedforward and recurrent connections (Fig. 4A). First, feedforward input from both juice channels is excitatory to one node and inhibitory to another node, enabling overlapping value representations. However, the strengths of these connections depend on juice type – to maintain equivalent value representations for the two juices, the connection from Juice A must be stronger than that from Juice B for the same quantity. Based on the functional roles, we refer to the nodes receiving excitatory and inhibitory feedforward inputs as the sensory and decision nodes, respectively.

Second, the recurrent connections support the dynamic evolution of value representations (Fig. 4B,C). Each node receives strong self-excitation, supporting the maintenance of encoded values and choices during delay periods. In contrast, connections between the sensory and decision nodes are asymmetric. Feedforward excitation from the sensory to the decision node enables a dynamic representation of working memory – under asymmetric inputs, Offer1 first activates the sensory node and then propagates to the decision node (Fig. 4C). Strong self-recurrent excitation prolongs the effective time constant of the circuit, slowing this propagation and producing rotational trajectories in the state space.

Feedback inhibition from the decision to the sensory node supports value comparison. In the reduced circuit, decision node activity at the end of Phase 1 reflects the Offer1 value and suppresses the sensory node (Fig. 4C, right). In Phase 2, the sensory node becomes active only if the Offer 2 input overcomes this suppression. With Offer 2 fixed at 4B, reactivation occurs when Offer 1 is 0A, 1A, or 2A, but not after 3A or 4A. Thus, sensory node activity in Phase 2 distinguishes trials according to the relative offer values. Because decision node activity eventually follows sensory node activity, the network shows lower activity for choosing Offer 1, and higher activity for choosing Offer 2. The full RNN showed a similar dependence of Offer 2 encoding on Offer 1 values. The sensory direction identified by latent-circuit inference was similar to the initial Offer 1 value-encoding direction (Fig. 3), and the response along this direction to the same second offer was more strongly suppressed by larger Offer 1 values (Fig. 4C, left).

Dynamic evolution during the delay period is evident in vector fields of the inferred circuit, similar to those of the full RNN (Fig. 4B). We further examined how network parameters shape this transformation through eigenvector decomposition of the recurrent connections (Fig. S4). Across RNN ensembles, the real part of eigenvalues nearly cancels intrinsic leakage, yielding spiral dynamics without net convergence or divergence. The imaginary part determining the rotational speed also remains consistent across RNN ensembles, with trajectories rotating about 90 degrees over the 500 ms delay. Despite the correspondence between rotation speed and delay durations, the solution identified by the RNNs remains robust when trained with varying inter-offer periods (Fig. S5). However, varying the delay period between Offer 2 and choice-to-action mapping distorted the choice output, creating a bias toward Offer 1 with longer second delays. This bias reflects the decay of activity toward the rotation center during the prolonged delays, shifting the population state toward the region associated with Offer 1 choices.

### Order-based choice signatures in experimental data and RNN

Analysis of RNN activity and latent circuit inference suggested a potential mechanism underlying order-based choices. To compare its neural signatures with experimental data, we examined the relationship between the encoding of Offers 1 and 2 using the bivariate regression analysis previously applied to sequential-stimuli perceptual (Romo and Salinas, 2003) and economic decision-making (Blanchard et al., 2015; Strait et al., 2015; Azab and Hayden, 2017; Ballesta and Padoa-Schioppa, 2019). For each Phase *j* (where *j* = 1 or 2), the activity of unit 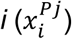 was modeled as:

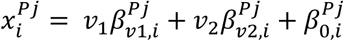

where *v*_1_ and *v*_2_ represent the values of Offer 1 and Offer 2, and *β* terms denote regression weights. To avoid spurious relationships arising from correlations between offer values, we used a test set of offer pairs with no correlation between *v*_1_ and *v*_2_ (Methods).

We then examined the relationships among offer-value tuning coefficients and order-choice preference in the order-task RNN (Fig. 5A-D) and the monkey data (Fig. 5E-L), which exhibited similar qualitative patterns. First, Offer 1 value tuning in Phase 1, 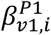 , was negatively correlated with order-choice preference (AUC_order)_ in both the RNN (Fig. 5A) and monkey data (Fig. 5E,I). As noted for Fig. 2, this correlation can arise from overlapping current-offer value representations across phases and the behavioral relationship between value and choice. Consistently, Offer 1 tuning in Phase 1, 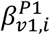 , was positively correlated with Offer 2 tuning in Phase 2, 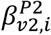 (Fig. 5B,F,J).

Second, during Phase 2, the memorized value of Offer 1 was encoded oppositely to the incoming value of Offer 2, with a negative correlation between 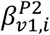 and 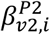 (Fig. 5C,G,K). These tuning relationships are consistent with previous findings in similar sequential-offer choice tasks (Strait et al., 2015; Azab and Hayden, 2017). In the RNN, the opposing Phase 2 tuning results from inhibition from the decision node to the sensory node in the inferred latent circuit (Fig. 4A). In the monkey data, on the other hand, this pattern does not uniquely identify that mechanism. When binary order choice was included as an additional regressor in Phase 2, the negative correlation between the two offer-value coefficients did not differ significantly from the permutation-based null distribution (Fig. S6). However, this does not establish that categorical choice fully accounts for the original correlation. Rather, opposing Phase 2 tuning could reflect graded relative value coding, categorical order-choice-related activity, or both, arising locally or through interactions with other regions (see Discussion).

By contrast, previous studies have reported more variable relationships between the initial encoding and later memory presentation of Offer 1, including both positive (Azab and Hayden, 2017) and negative correlations (Cavanagh et al., 2018). In our data, the correlation between 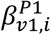 and 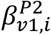 was negative or weak — near zero in one monkey (Fig. 5D,H,L). A similar attenuation also appeared in some order-task RNNs (Fig. S7). In the model, this effect arises from the dynamic transformation of the Offer 1 representation across phases, which altered the regression vector 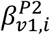 , and weakened its negative correlation with 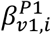.

To connect these tuning relationships to the population-level geometry observed in RNNs, we further examined value-encoding directions across successive task epochs in OFC pseudo-populations and order-task RNNs (Fig. 6). Cosine similarities revealed progressive changes of Offer 1 encoding, alignment of current-offer value representations across phases, and an opponent organization of Phase-2 offer-value directions in both OFC and RNNs (Fig. 6C,D). However, their magnitudes and timing of these relationships differed: OFC Phase 2 patterns appeared to combine features observed during Offer 2 presentation and the subsequent delay in order-task RNNs (e.g., SO_1-_SO_2 a_nd SD_1-_SD_2 i_n RNNs vs. SO_1-_SO_2 i_n OFC). This temporal discrepancy was reduced in RNNs trained on the monkey task, which lacked a second delay, suggesting that task structure may contribute to the observed differences (see Discussion).

### Comparison to identity-based choice in attractor dynamics

So far, we have focused on order-based choice in sequential decision-making, with dynamics that are largely independent of juice identity. However, identity-based choice representations have been found to be important in economic choice tasks where option identity remains relevant to behavior (Hunt et al., 2012; Chau et al., 2014; Rustichini and Padoa-Schioppa, 2015). To compare mechanisms across choice reference frames, we modified the choice-to-action mapping so that left/right actions were associated with juice identity rather than presentation order (Fig. 7A), following an experimental paradigm in which juice-based choice signals were observed (Ballesta and Padoa-Schioppa, 2019). Choice decoding confirmed juice-based choice signals (Fig. 7B). We refer to these networks as juice-task RNNs.

As in the order-task RNNs, activity during Phases 1 and 2 was largely confined to two dimensions, accounting for 91.6 % of the variance in an example juice-task RNN. However, unlike the order-task RNNs, these networks exhibited two distinct encoding axes during Phase 1, corresponding to the values of Juice A and B (Fig. 7C, left). These axes formed an “L” shape, meeting at the origin, with offer values encoded and maintained monotonically along each axis. During the Offer 1 presentation, activity magnitude increased while remaining aligned to the corresponding axis. During the inter-offer period, the representation of Offer 1 remained largely stable (Video 3). Notably, these axes were subsequently repurposed to represent choice signals (Fig. 7C, right): following Offer 2, choice representations aligned with the same axes used for encoding and maintaining offer values (Video 4).

Stable working memory and choice representation in the juice task are reminiscent of attractor dynamics, where each juice axis acts as a distinct attractor, supporting both working memory and decision-making. Consistently, vector field analysis during the delay period revealed attraction toward the two axes, resembling line attractors for each axis (Fig. 7D). The inferred latent circuit consisted of two competing nodes, each receiving strong input from one juice and corresponding to its encoding axis in the population analysis. Strong recurrent connections provided positive feedback within each node, whereas moderate mutual inhibition between the nodes supported competition and choice. Additionally, each node received inhibition from the opposite juice input. The latent circuit’s dynamics recapitulated the RNN dynamics, with each node tuned to Offer 1 value in Phase 1, and to juice-based choice in Phase 2 for the corresponding juice.

Analysis of value representations in juice-task RNNs showed a positive correlation between 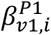 and 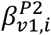 (Fig. 8C), reflecting stable value representations for each juice – a key distinction from the order task. Another feature is the discrepancy between correlation patterns in separate AB and BA trials versus pooled averages (Fig. 8A-D). Specifically, correlations between 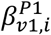 and 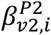 were positive when AB and BA trials were pooled, but negative when analyzed separately (Figs. 8A versus D). The latter reflects competition between the two value representations in juice-task RNNs, whereas averaging *β* terms across AB and BA trials mixes activity along distinct juice axes. This mixing broadens fluctuations in correlation patterns across RNNs, which can obscure encoding and choice schemes in juice-task RNNs. Conversely, the negative correlation between 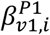 and 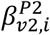 in the segregated trials contrasts with the juice- and phase-agnostic encoding of incoming offer values in the order-task RNNs, which may further help differentiate among choice reference frames.

These contrasting representational and dynamical features prompted us to examine whether the two coding schemes could coexist within the same network. Some networks exhibited both juice-based and order-based choice representations (Fig. 1H). These networks tended to show higher-dimensional activity during the delay period than those exhibiting either representation alone (Fig. 9). They also combined the dynamical features of both coding schemes: juice-based representations were associated with relatively stable, attractor-like dynamics, whereas order-based representations underwent rotation-like transformations. Thus, order-based rotational dynamics are not restricted to juice-agnostic networks but can coexist with juice-based stability within the same network.

## Discussion

Here, we found that in monkey OFC, order-based choice signals emerge rapidly and become dominant during sequential-offer economic choice when behavior reporting was not restricted to juice identity. This contrasts with the juice-based choice signals reported in the previous studies using identity-based choice-to-action mappings (Ballesta and Padoa-Schioppa, 2019). RNNs constrained to reproduce these signals suggested a mechanism for working memory and decision formation that differs from the classical picture of competition between populations that stably preserve each option’s identity. Instead, the model relied on asymmetric feedforward and recurrent interactions, which generated overlapping representations of the currently attended offer, largely independent of offer identity, and transformed this activity into a memory representation that could be compared with the subsequent offer. Importantly, this mechanistic account also yielded relationships between offer-value coding and choice preference, which were qualitatively consistent with those observed in monkey OFC.

A central feature of this mechanism is the transformation of value representations across task phases. Such reformatting provides a solution to a core problem in sequential choice: how the brain can distinguish the currently viewed offer from the remembered one while still comparing them on a common value scale. Similar regression-based analyses and related subspace transformations affecting firing covariance structures have been reported in other sequential-offer value-based decision-making tasks, including gambling paradigms (Blanchard et al., 2015; Strait et al., 2015; Azab and Hayden, 2017; Yoo and Hayden, 2020; Johnston et al., 2024).

Conceptually, the mechanism of order-based choices aligns with normative proposals that choices under sequential offers are implemented through shifts of attention, such that a common neural pool evaluates the currently attended offer while the previously presented offer serves as a comparison threshold in an accept-reject computation (Kacelnik et al., 2011; Hayden, 2018; Hayden and Moreno-Bote, 2018).

Similar sensory-to-memory transformations have also been proposed in working memory and perceptual decision-making based on asymmetric recurrent connections between sensory and memory nodes (Miller and Wang, 2006). Our work further suggests that economic choice may utilize a similar circuit motif with inputs extended to multiple channels and their connections modified to convert juice quantities into value representations. Notably, the representational transformations described in earlier works do not require rotational dynamics per se. Although our RNN analyses and inferred recurrent structure are consistent with a gradual, rotational evolution during the delay period, the essential computation supported by the neural signatures is the conversion of current value information into a memory representation that supports later comparison. Future studies need to investigate the specific temporal profile of this transformation and what advantage it provides.

Because our original RNN task postponed and randomized the mapping from choice to action, it allowed us to isolate abstract order-based decision formation from spatial response planning, as in previous juice-based paradigms (Cai and Padoa-Schioppa, 2014, 2019; Ballesta and Padoa-Schioppa, 2019). In tasks without this separation, the relative recruitment of abstract and action-based reference frames can depend on task structure (Hunt et al., 2013). To test whether the relevant order-based signatures depended on delaying the mapping, we trained additional RNNs using the monkey task structure, with spatial information available during offer presentation (Fig. 10). These networks exhibited a transition from order-based to spatial-based choice and reproduced key representational features observed in the original networks and monkey recordings: transformation of the Offer 1 value representation during the delay, alignment of current-offer value representations across Phases 1 and 2, and an opponent organization of the Phase-2 offer-value coefficients. Thus, these signatures also emerged when spatial information was available during offer presentation, indicating that they were not specific to the delayed-mapping task.

These results support the relevance of studying order-based representations even when the task permits spatial response planning. However, the monkey-task RNNs represented spatial information from the beginning, exhibited higher-dimensional population activity, and showed less temporal segregation between order-based and spatial-based choices. These findings suggest that order-based representations can coexist with spatial processing, rather than necessarily constituting a completed abstract decision stage before action selection begins. The delayed-mapping task therefore provides a useful setting for isolating the order-based component of the computation, but does not capture all interactions present when spatial information is available earlier. Also, the comparison to data was limited to the pseudo-population analysis characterizing task-related representational geometry rather than the trial-by-trial evolution of a simultaneously recorded population state. Whether order-based value encoding and feedback inhibition identified in our original RNNs are modified or complemented by spatial computations needs to be investigated. This would also require comparison to larger simultaneously recorded populations and finer temporal analyses that could link population-state fluctuations to trial-by-trial choices and resolve potentially rapid transitions from graded value comparison to categorical choice.

Previous theoretical frameworks propose that decisions emerge through distributed interactions among abstract and action-related representations (Cisek, 2012; Hunt and Hayden, 2017). OFC recordings have revealed choice-related value dynamics (Rich and Wallis, 2016; McGinty and Lupkin, 2023). Related studies have implicated vmPFC in comparing integrated offer values (Strait et al., 2014) and dorsal ACC (Azab and Hayden, 2017). Moreover, simultaneous OFC-ACC recordings linked OFC value dynamics to choice-related ramping in ACC (Kennerley et al., 2011; Hunt et al., 2015; Balewski et al., 2023), while lateral prefrontal cortex implicated this region in linking values and choices to actions (Cai and Padoa-Schioppa, 2014; Hunt et al., 2018). Our model complements these findings by describing how value representations can be maintained and transformed to support order-based comparison. Although comparison occurs within the modeled circuit, the recordings do not distinguish among potentially complementary OFC contributions: performing comparison locally, providing transformed inputs to comparison elsewhere, or receiving comparison-related feedback. Multi-area recordings spanning OFC, vmPFC, ACC, and lateral PFC, combined with models incorporating region-specific inputs, outputs, and interareal connectivity, could clarify how local and distributed computations support the coexistence and transformation of multiple-choice reference frames.

More broadly, our study supports a general view of cognition in which recurrent circuits adapt their dynamics to the computational demands of the task. Across working memory and related domains, RNN models have shown that changes in delay structure, memory-manipulation requirements, robustness constraints, or temporal multiplexing can favor persistent attractors, feedforward propagation, activity-silent memory, or oscillatory coding (Masse et al., 2019; Orhan and Ma, 2019; Kozachkov et al., 2020; Zhou et al., 2023). Our results extend this principle to economic decision-making where choices involve multiple attributes.

Complementing attractor-based accounts of stable, identity-specific representations, our model shows how order-based choice signals can arise through dynamical reorganization of value representations across task phases. In a subset of networks, order-based rotations coexisted with attractor-like stability of juice-based representations. This coexistence offers a possible framework for relating diverse neural signatures across value-based choice tasks. How task demands influence the relative engagement of these computations – potentially through context-dependent gating or input alignment with specific subspaces – remains to be investigated. Overall, our results underscore the flexibility of recurrent computations and support the view of decision-making as a task-dependent process spanning multiple representational formats.

## Author Contributions

J.G., and S.L designed and performed RNN research; B.V. performed reinforcement learning–based RNN simulations; K.N., and X.C. designed and performed animal research; J.G., K.N., X.C., and S.L. wrote the paper.

## Acknowledgments

We thank John Rinzel, Jeffrey Erlich, and Alex Reyes for their constructive suggestions throughout the research and for providing feedback on the manuscript. We thank Yuqing Su for contributing to the training of RNNs on the monkey task. J.G. and S. L. were supported by STI2030-Major 38 Projects, No.2021ZD0203700/2021ZD0203705 and NYU Shanghai Boost Fund. K.N. and X.C. were supported by STI2030-Major 38 Projects, No.2021ZD0203700/2021ZD0203702 and NYU Shanghai Boost Fund. X. C. and S.L. acknowledge the support of the NYU-ECNU Institute of Brain and Cognitive Science at NYU Shanghai S. L. also acknowledges the support of the NYU Shanghai Center for Data Science.

## Supplementary Figures

**Supplementary Figure S1.**
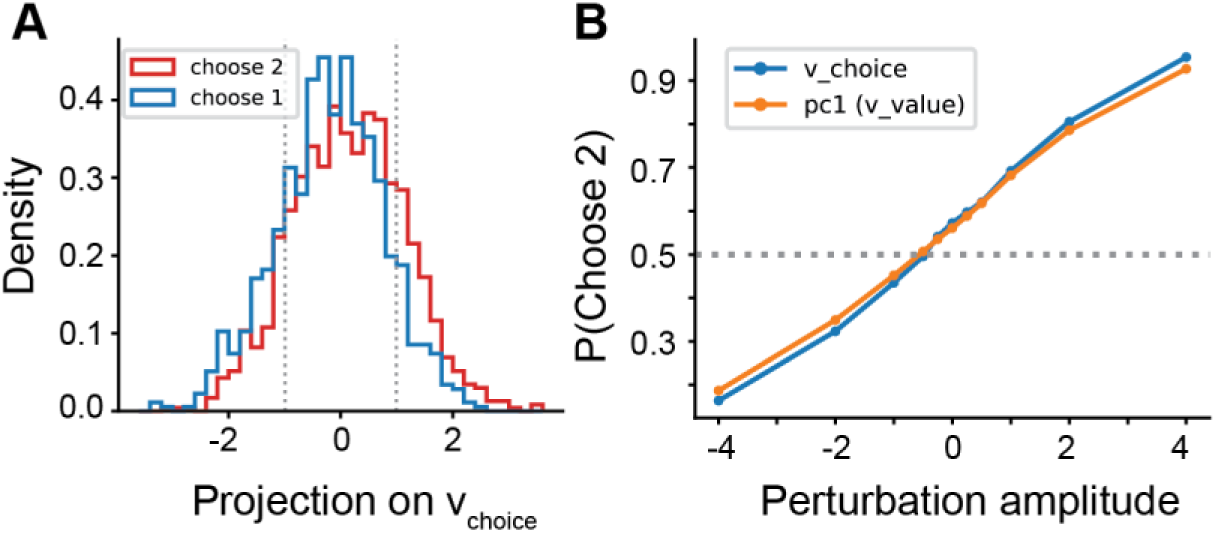
Effect of pre-Offer2 activity fluctuations on choice in an example order-task RNN. (A) Distributions of activity fluctuations projected onto a linear choice-predictive axis, v_choice,_ grouped by subsequent choice (Offer 1, blue; Offer 2, red). Analyses were restricted to offer conditions where both choices occurred: (q_A,_q_B)_ in {(1,1),(1,2),(2,3),(2,4),(3,5),(3,6),(4,8)}. v_choice w_as estimated from trial-by-trial deviations from condition-mean population activity immediately before Offer 2 onset. (B) Probability of choosing Offer 2 following perturbations along v_choice (_blue) or the first principal component (∼v_value,_ orange). The two axes were nearly aligned (cosine similarity, 0.988), and perturbations along them produced similar changes in choice probability.

**Supplementary Figure S2.**
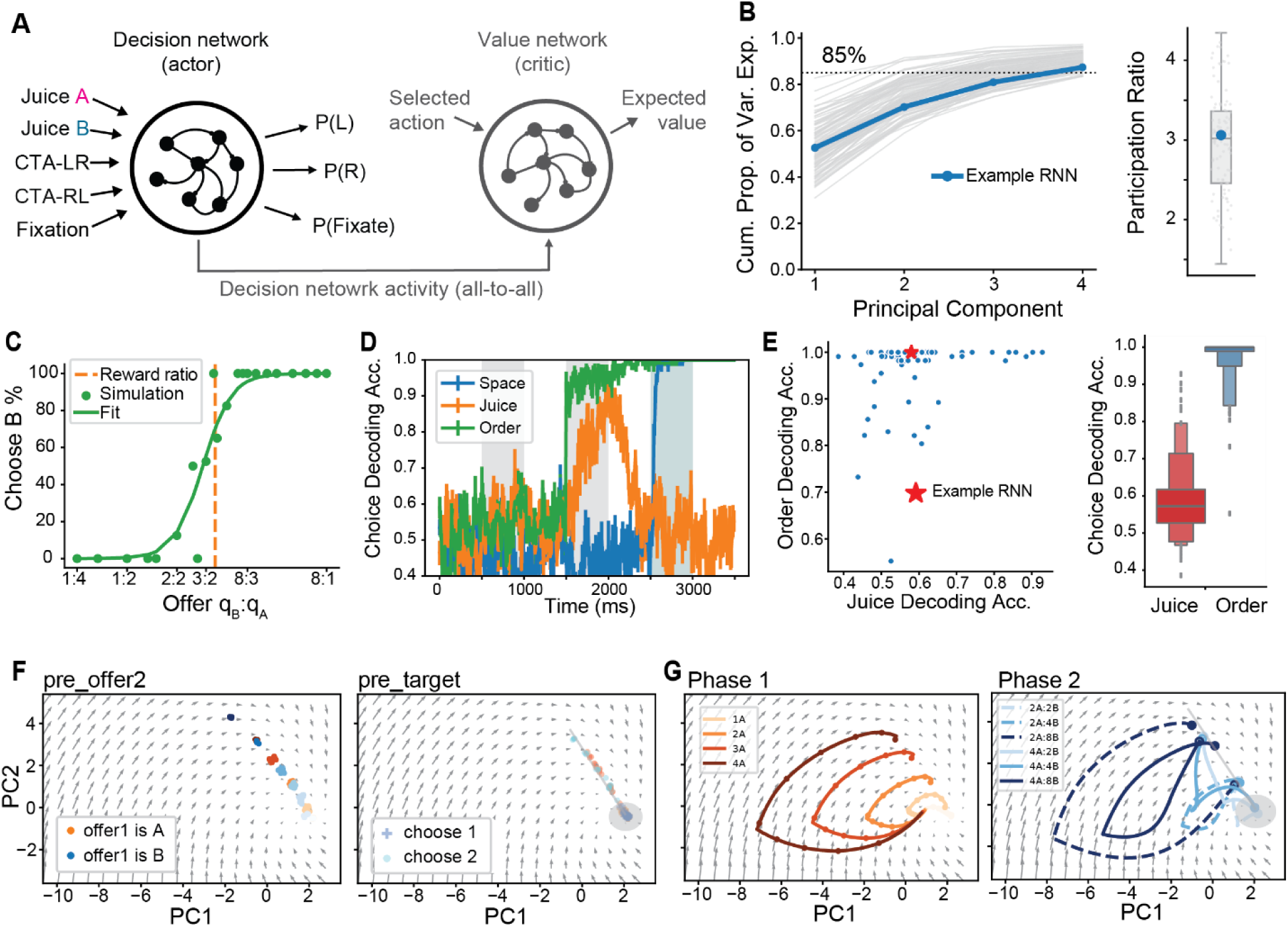
Order-task RNNs trained using model-free reinforcement learning. (A) Actor–critic architecture. The decision (actor) network had the same architecture and inputs as the supervised RNNs, while an auxiliary value (critic) network estimated expected reward to provide a baseline for REINFORCE training. All analyses focus on the actor networks. (B) Cumulative variance explained by principal components (left) and participation ratios (right) across RL-RNNs. (C) Psychometric curve of the example network. (D) Choice-decoding accuracy for the example network. (E) Juice- and order-based choice-decoding accuracy across RL-RNNs, shown as a scatter plot (left) and distributions (right). The red star identifies the example network. (F, G) Population activity and trajectories, projected onto the first two principal components. The format is the same as in Fig. 3. The example network exhibited projected dynamics qualitatively similar to those of supervised RNNs, despite the higher dimensionality and potentially greater diversity of the RL-RNN ensemble.

**Supplementary Figure S3.**
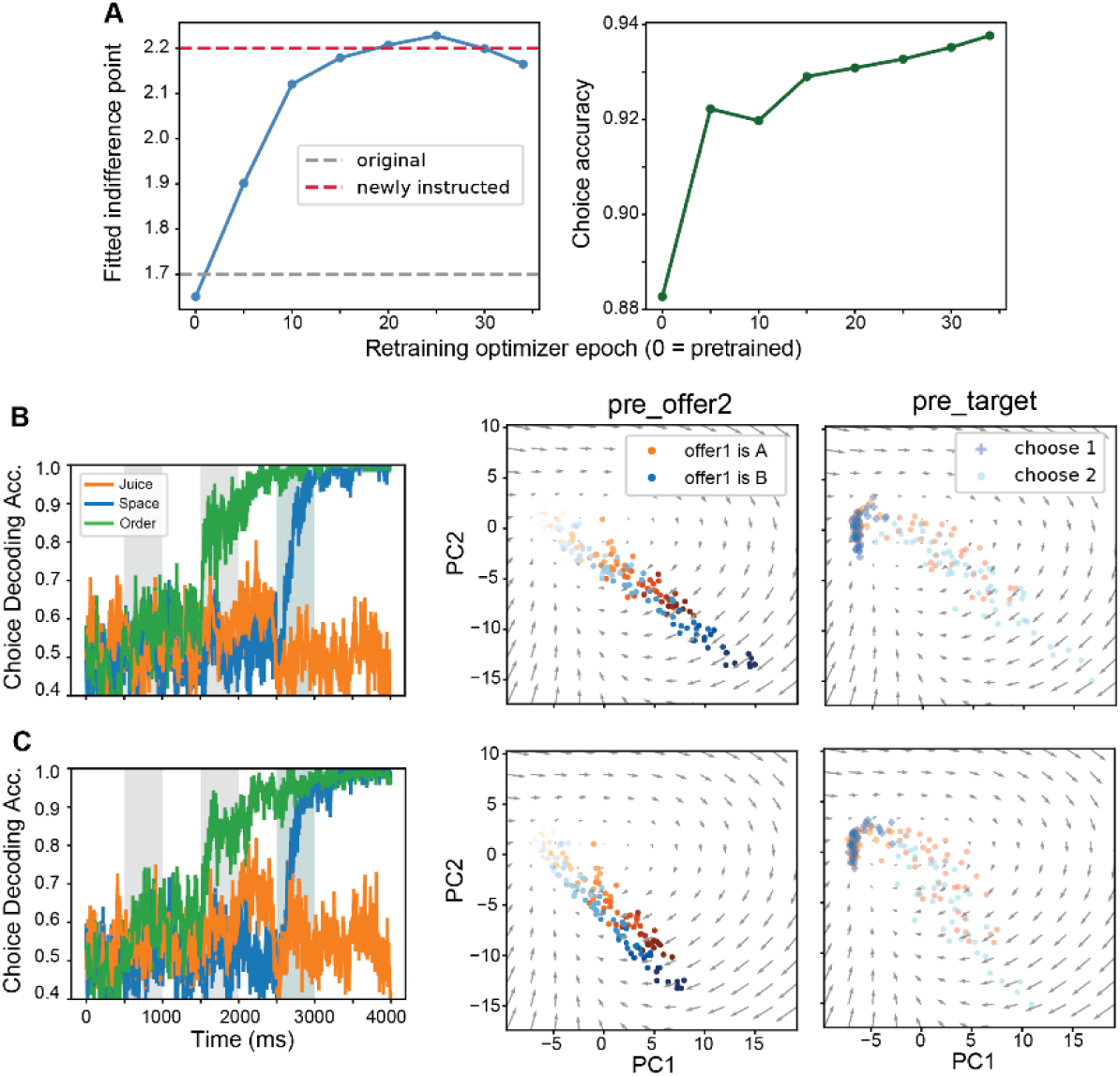
Effects of changing the instructed indifference point on RNN representations and dynamics. (A) Fitted indifference point (left) and choice accuracy (right) during retraining of an example order-task RNN. (B,C) Choice decoding (left) and population activity projected onto the first two PCs at the end of Phase 1 (middle) and Phase 2 (right), before (B; same example as in the main figures) and after (C) retraining. Subtle changes in the weight matrices rescaled the value representation for the two juices in the low-dimensional PCs, while the overall dynamics are largely preserved.

**Supplementary Figure S4.**
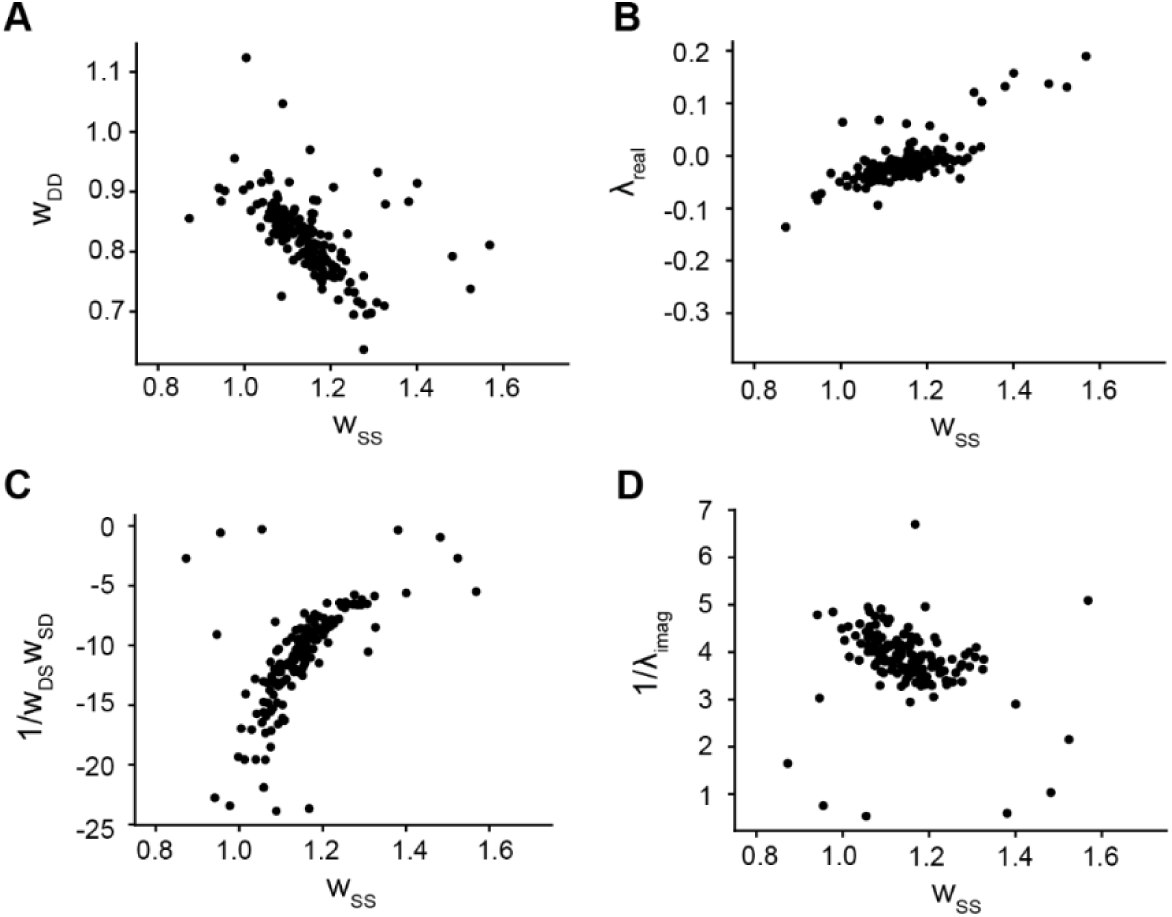
Parameter dependence of the reduced circuit. (A,B) Relationship between recurrent excitation of sensory node *w_SS_* and decision node *w_DD_*, and their connection to the eigenvalues of the recurrent dynamics. Empirically, the sum of recurrent excitation remains nearly constant, resulting in a real eigenvalue component given by 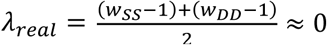 , and almost perfect cancellation of intrinsic decay during the delay period. (C,D) Relationship between recurrent excitation *w_SS_* and recurrent inhibition generating a negative feedback loop *w_DS_w_SD_* and their connection to eigenvalues. The imaginary eigenvalue component λ 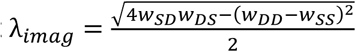 is also nearly constant around 1/4. Given an intrinsic RNN time constant of *τ* =100ms, the oscillation period 2πτ/λ*_imag_* is approximately 1800-3000ms, corresponding to 1/4 cycle of oscillation in the vector fields, and aligning with the delay period time scale.

**Supplementary Figure S5.**
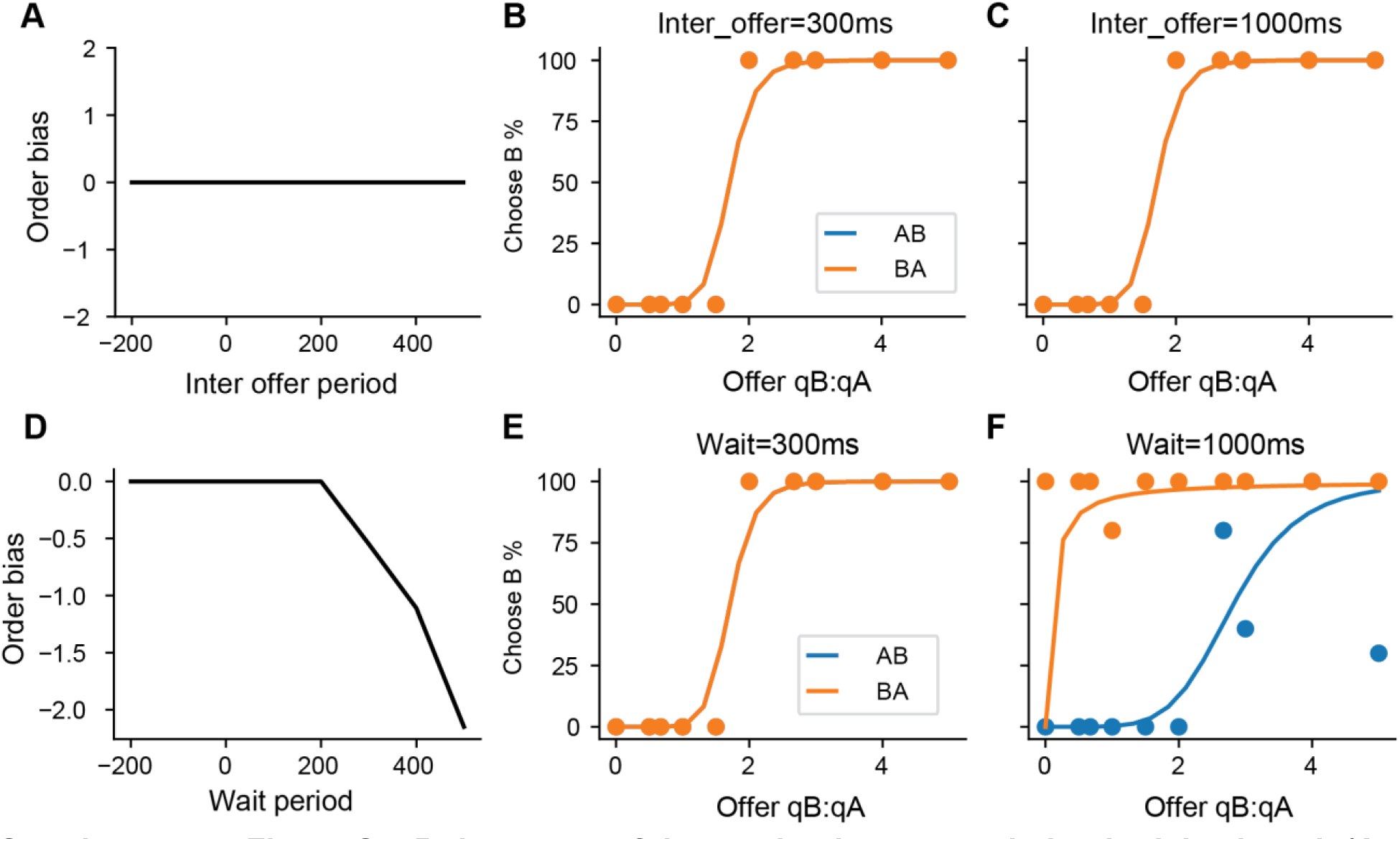
Robustness of the mechanisms to variation in delay length (A-C) and wait period (D-F). A separate set of RNNs was trained with variable inter-offer duration and variable wait period (300ms to 1000ms). (A) Order bias remained unchanged across the trained range of inter-offer durations. (B,C) Psychometric curves from example RNNs tested at the shortest and longest inter-offer durations, showing overlap between AB and BA trials. (D-F) When RNNs were trained with variable wait periods, order bias emerges at longer wait periods (same layout as in A-C).

**Supplementary Figure 6.**
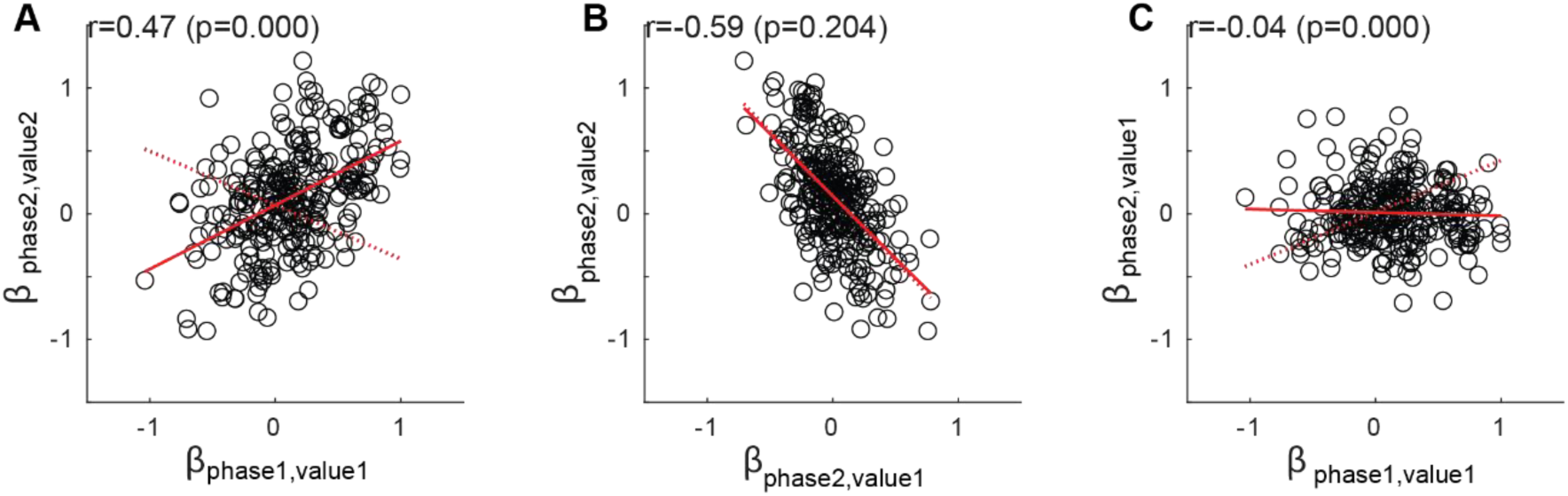
Relationships among offer-value tuning coefficients after controlling for binary choice. (A) Offer 1 tuning in Phase 1 vs. Offer 2 tuning in Phase 2. (B) Offer 1 versus Offer 2 tuning in Phase 2. (C) Offer 1 tuning in Phase 1 versus Phase 2 (n = 288 neurons). Phase 2 activity was regressed jointly on Offer 1 value, Offer 2 value, and binary order choice. Regressions were performed separately for each time window and each of four trial sets, using trial-type averages of z-scored firing rates and predictors. Regression coefficients were averaged across trial sets, and pairwise correlations were computed across neurons passing ANOVA filtering (p < 0.001). Statistical significance was assessed using permutation tests in which trial-by-trial firing rates were shuffled within each trial set, and regressions and coefficient correlations were recomputed (1,000 permutations). Solid red lines indicate observed linear fits, and dashed red lines indicate mean slopes from the permutation-based null distributions. Reported p-values are permutation-based.

**Supplementary Figure S7.**
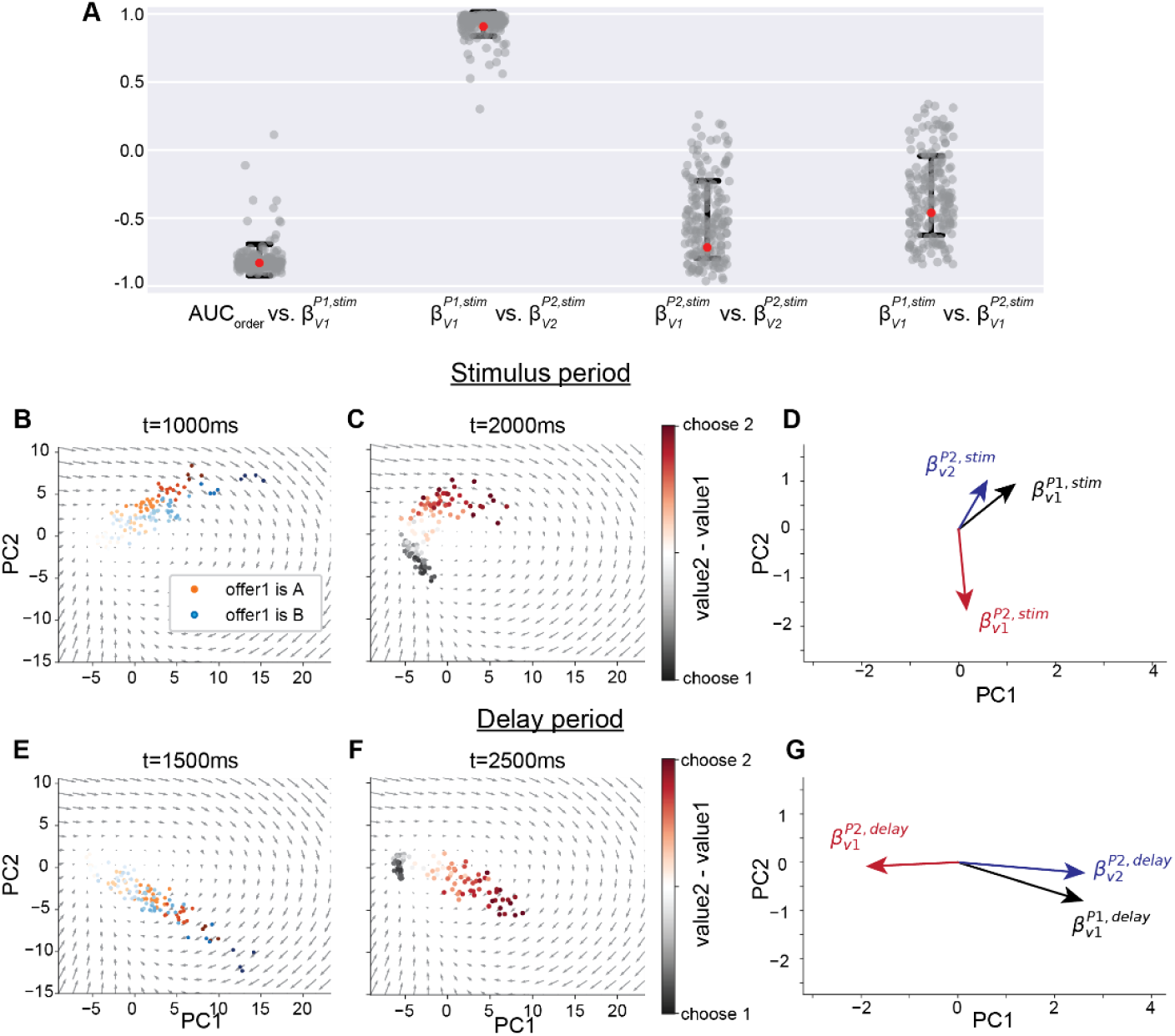
Distribution of correlation across RNNs and evolution of regression vectors and their impact on correlation patterns. (A) Correlation coefficients across RNN ensembles, obtained from time-averaged activities at the end of the stimulus period. Results from example RNNs in Figures 3A-D were marked by red dots. (B-G) Correlation patterns during the stimulus period (B-D) and the delay period (E-G). (B-D) Population activity (same as in Figures 3A-D) at the end of the stimulus period in Phase 1 (B) and Phase 2 (C), and corresponding regression coefficient vectors 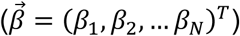 projected onto the PC plane (D). In Phase 1, activity aligns in a single direction regardless of offer identity and is consistent with the regression vector 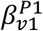 (B; black in D). By contrast, Phase 2 activity separates into two clusters corresponding to choice (C; choosing Offer 1: black; Offer 2: red). Here, 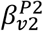 aligns with Offer 2 choices (blue in D), while 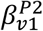 (red in D) is a combination of Offer 1-choice activity (black in C) and the opposite of Offer 2-choice activity (red in C). (E-G) Same format as (B-D), but showing activity and regression vectors at the end of the delay period. Although activity evolves through rotation dynamics over time, the relationship between activity and regression vectors remains the same, with 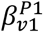 and 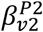 rotating in parallel with population activity (G). In contrast, 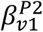 changes more drastically because Offer 1-choice activity decays, leaving it nearly opposite to Offer 2-choice activity. Since the correlations between *β*’s reflect the cosine of the angle between 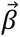 vectors, the negative correlation between 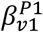 and 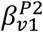 is attenuated during the stimulus period compared to the delay period. This occurs because, during the stimulus period, the orthogonal transformation of Offer 1 representations across phases (black in C) strongly influences the regression vector, 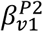.

## Video Legends

**Video 1.**
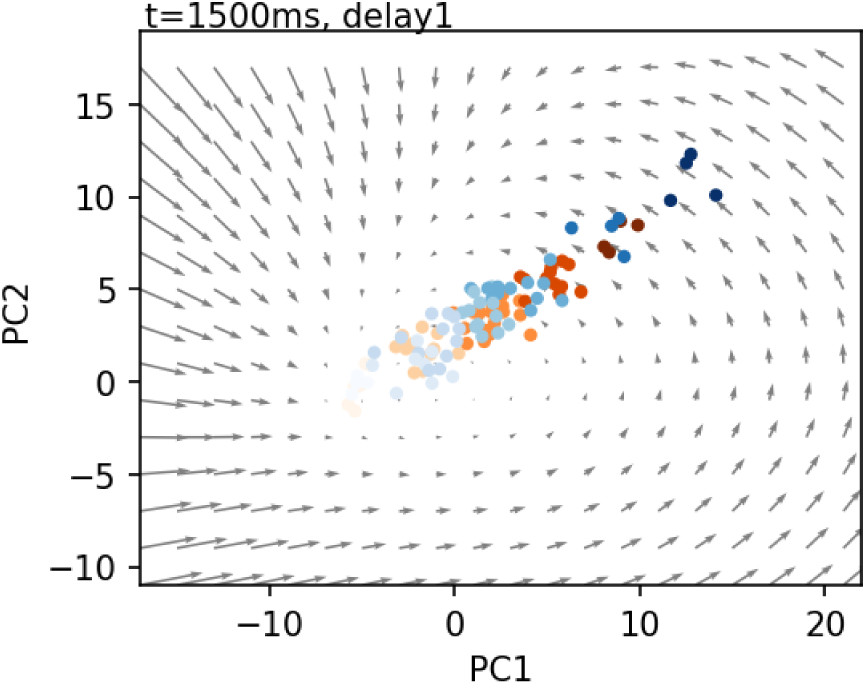
Dynamics of population activity during Offer 1 encoding and maintenance in the order-task RNN. Progression of activity in an example RNN during Phase 1, projected onto the first two principal components (PCs). Offer 1 was presented from 500 to 1000 ms (orange: Juice A; blue: Juice B; darker shades indicate larger Offer 1 quantities), followed by a delay period from 1000 to 1500 ms. Activity initially increased proportionally to the Offer 1 value in a juice-agnostic manner and then rotated during the delay period. Gray arrows indicate the vector field of the RNN dynamics in the absence of input. Figure 3A shows a snapshot of the activity at the end of Phase 1, with PC 2 flipped.

**Video 2.**
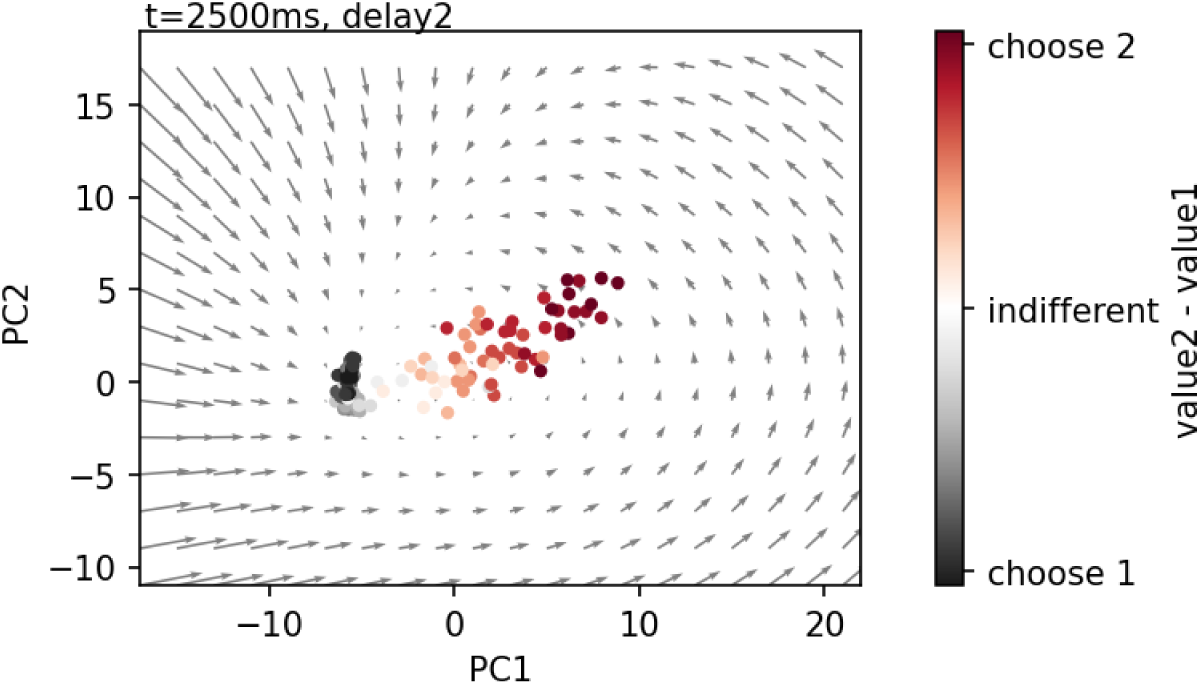
Dynamics of population activity during value comparison and choice in the order-task RNN. Progression of activity in an example RNN during Phase 2, projected onto the first two principal components (PCs). Offer 2 was presented from 1500 to 2000 ms, followed by a delay from 2000 to 2500 ms. Color indicates order-based choice: black for Offer 1 and red for Offer 2, with shading representing the value difference of value difference between Offer 1 and Offer 2. Toward the end of the delay, trajectories were nearly aligned along a common choice axis, with Offer 1-choice activity located near the center of rotation and Offer 2-choice activity broadly distributed. Gray arrows indicate the vector field of the RNN dynamics in the absence of input. Figure 3B shows a snapshot of the activity at the end of Phase 2, with PC 2 flipped.

**Video 3.**
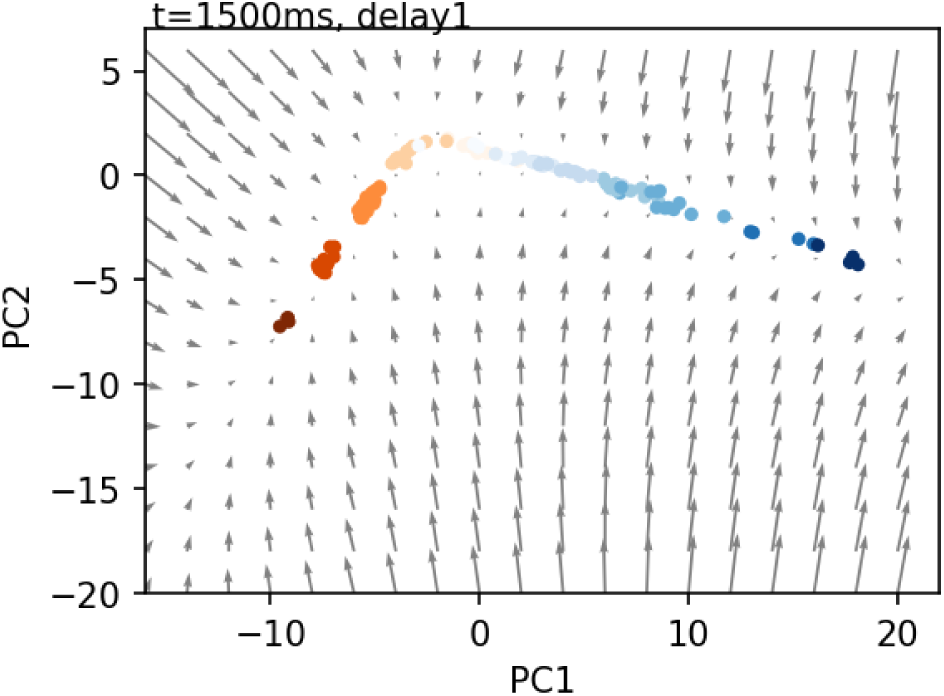
Dynamics of population activity during Offer 1 encoding and maintenance in the juice-task RNN. Progression of activity in an example RNN during Phase 1, projected onto the first two principal components (PCs). Offer 1 was presented from 500–1000 ms (orange: Juice A; blue: Juice B), followed by a delay period from 1000–1500 ms. Activity evolved along juice-specific axes and remained stable during the delay.

**Video 4.**
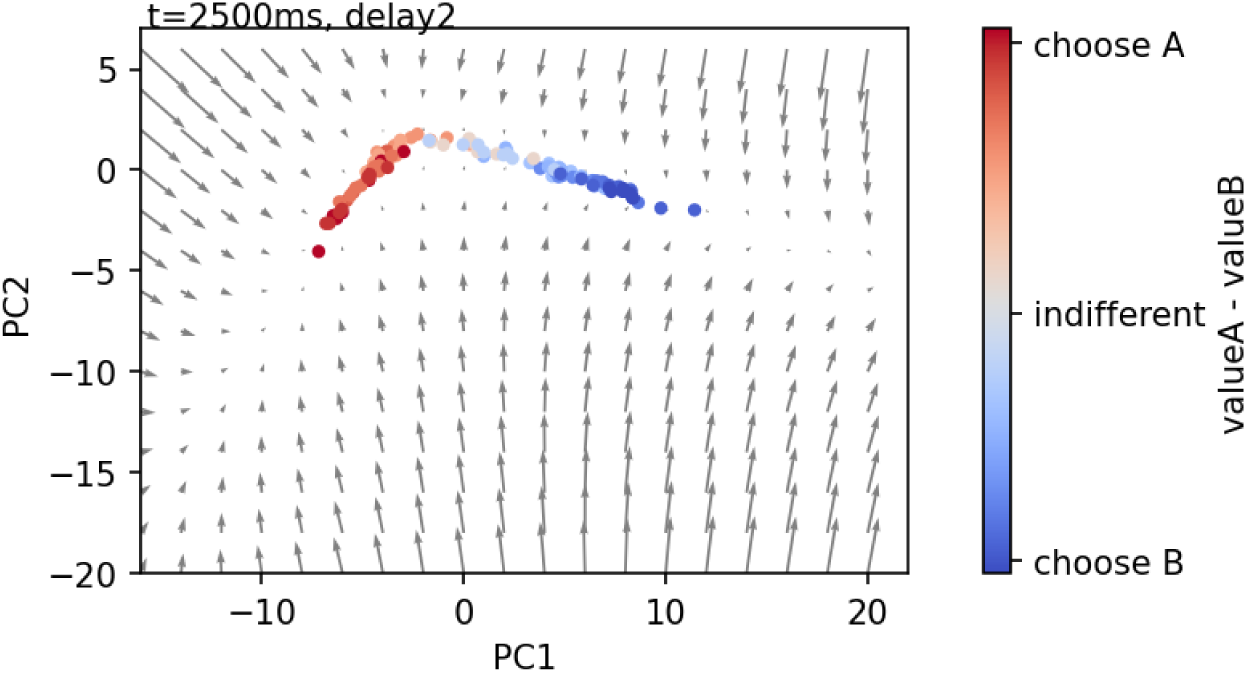
Dynamics of population activity during value comparison and choice in the juice-task RNN. Progression of activity in an example RNN during Phase 2, projected onto the first two principal components (PCs). Offer 2 was presented from 1500–2000 ms, followed by a delay from 2000–2500 ms. During the delay, choice-related activity aligned with the juice-encoding axes (red: choose Juice A; blue: choose Juice B), and the distance from the origin reflected the value difference between the two offers (represented by a color gradient).

